# Diving into broad-scale and high-resolution population genomics to decipher drivers of structure and climatic vulnerability in a marine invertebrate

**DOI:** 10.1101/2024.01.29.577252

**Authors:** Audrey Bourret, Christelle Leung, Gregory N. Puncher, Nicolas Le Corre, David Deslauriers, Katherine Skanes, Hugo Bourdages, Manon Cassista-Da Ros, Wojciech Walkusz, Nicholas W. Jeffery, Ryan R. E. Stanley, Geneviève J. Parent

## Abstract

Species with widespread distributions play a crucial role in our understanding of climate change impacts on population structure. In marine species, population structure is often governed by both high connectivity potential and selection across strong environmental gradients. Despite the complexity of factors influencing marine populations, studying species with broad distribution can provide valuable insights into the relative importance of these factors and the consequences of climate-induced alterations across environmental gradients. We used the northern shrimp *Pandalus borealis* and its wide latitudinal distribution to identify current drivers of population structure and predict the species vulnerability to climate change. Individuals sampled across 24° latitude were genotyped at high geographic-(54 stations) and genetic-(14,331 SNPs) resolutions to assess genetic variation and environmental correlations. Four populations were identified in addition to finer substructure associated to local adaptation. Geographic patterns of neutral population structure reflected predominant oceanographic currents, while a significant proportion of the genetic variation was associated with gradients in salinity and temperature. Adaptive landscapes generated using climate projections suggest a larger genomic offset in the southern extent of the *P. borealis* range, where shrimp had the largest adaptive standing genetic variation. Our genomic results combined with recent observations point to the non-recovery in southern regions and an impending vulnerable status in the regions at higher latitude for *P. borealis*. They also provide rare insights into the drivers of population structure and climatic vulnerability of a widespread meroplanktonic species, which is crucial to understand future challenges associated with invertebrates essential to ecosystem functioning.

## Introduction

Marine species often exhibit substantial abundance and extensive geographic ranges, covering a spectrum of environmental conditions with few conspicuous barriers to dispersal (Mach et al. 2011; Stanley et al. 2018; Knutsen et al. 2022). These attributes render marine species ideal candidates for investigating the underlying factors – both neutral and adaptive – that shape population structure (Nielsen et al. 2009). Two critical features that constrain population structure in many marine species include large-scale oceanographic currents, which homogenize genetic structure by facilitating gene flow, and large population sizes, which minimizes the effects of genetic drift. Still, environmental gradients can lead to local adaptation and genetic heterogeneity in the ocean, even under conditions of substantial migration and gene flow (Nielsen et al. 2009; Tigano and Friesen 2016). Genotype-environment association approaches have revealed candidate adaptive loci among many taxa (e.g., Benestan et al. 2016; Jeffery et al. 2018; Kess et al. 2021; Ferchaud et al. 2022), indicating instances of local adaptation (Savolainen et al. 2013; Rellstab et al. 2015). Thus, while populations might appear homogeneous when assessing neutral loci, the same may not hold true for adaptive regions of their genome (Conover et al. 2006; Gagnaire et al. 2015). Nonetheless, the mechanisms guiding population structure and the relative significance of neutral versus adaptive processes for many marine species remain unclear.

Utilizing genomic tools such as genotyping-by-sequencing and whole genome sequencing can aid in identifying and understanding the relative importance of evolutionarily neutral and adaptative processes that shape species genetic composition (Hohenlohe et al. 2019). These tools are being increasingly applied to refine our understanding of stock structure and for the conservation of species experiencing myriad threats (Flanagan et al. 2018; Barbosa et al. 2020; Formenti et al. 2022). Genomic variation within a species is often uneven across its distribution. This heterogeneity can stem from a combination of geographic variation in evolutionary history, barriers to gene flow, and fine-scale adaptation to both historical and contemporary environmental conditions. The relative contributions of adaptation and neutral evolutionary processes can be evaluated using population and landscape genomics (Luikart et al. 2003; Rellstab et al. 2015), while functional genomics (e.g., transcriptomics) can be useful to pinpoint mechanisms and their intraspecific variation involved in response to changing environmental conditions and to improve inference of local adaptation (Forester et al. 2018; Keagy et al. 2023).

Genomics can also be leveraged to better understand species responses to climate change (Franks and Hoffmann 2012; Razgour et al. 2019; Capblancq et al. 2020a; Waldvogel et al. 2020; Bernatchez et al. 2023). Anticipated impacts of global climate change on the marine environment are numerous and encompass increased temperatures, rising sea levels, changes in salinity, decreased pH levels, and an increasing frequency and severity of extreme weather events (Pörtner et al. 2019), all of which are likely to impact species resilience and distribution. These shifts in environmental conditions are projected to lead to biological changes rooted in physiological constraints, including species’ range shifts, invasions, extinctions, and modifications to ecosystem functioning (e.g., Stige and Kvile 2017; Jeffery et al. 2018; Stanley et al. 2018; Layton et al. 2021). Increasing temperatures, in particular, are viewed as a critical factor driving species range shifts and abundance declines now and into the future (Perry et al. 2005; Cheung et al. 2009). Changing environmental conditions will also exhibit spatial heterogeneity, implying that the magnitude of change will differ throughout a species’ distribution, influencing genetic composition and a variable genetic/adaptive landscape (Layton et al. 2021). The response to change in selection pressure is impacted by the amount of standing genetic variation already present in a population, which could ease adaptation to new conditions (Barrett and Schluter 2008; Bernatchez 2016). Integration of genomic and environmental data with climate predictions enables the assessment of genomic offset through the comparison of adaptive landscape predicted under current and future environmental conditions (Capblancq et al. 2020a; Rellstab et al. 2021; Layton and Bradbury 2022). Genomic offset can be viewed as the expected shift in genomic optimum, or similarly as the amount of genetic adaptation required for populations to endure changing environmental conditions. Populations with higher genomic offset are expected to have more difficulty coping with predicted changes (Bay et al. 2018).

Northern shrimp (*Pandalus borealis* Krøyer, 1838) are a commercially harvested species distributed across the Arctic and North Atlantic Oceans, occupying habitats with a wide range of temperatures (−1 to 12 °C), salinities (23.4 to 35.7 ppm), and depths (9 to 1400 m; Allen 1959; Shumway et al. 1985). Northern shrimp are protandric hermaphrodites, where females spawn once a year in late summer and carry their eggs for approximately eight months until they hatch in spring (Koeller et al. 2009). As a meroplanktonic species, northern shrimp have a biphasic life history, with a pelagic larval phase where larvae are found mainly in surface layers (Ouellet and Allard 2006) followed by a benthic juvenile and adult phases. This life history favours a high dispersal potential and gene flow during the pelagic larval phase, before recruiting to the benthic environment where dispersal is expected to be generally more limited (Shumway et al. 1985). Temperature plays a key role regulating mortality and the phenology of pelagic and benthic life history phases (Ouellet and Chabot 2005; Koeller et al. 2009; Daoud et al. 2010; Arnberg et al. 2013). Under laboratory conditions, larvae hatch earlier and develop faster at higher temperature, but eggs show reduced hatching success (Brillon et al. 2005; Ouellet and Chabot 2005; Daoud et al. 2010; Arnberg et al. 2013). Similarly, adults show higher metabolic rate but lower survival under elevated temperature (Chemel et al. 2020; Guscelli et al. 2023). The response to changing temperature, however, is not uniform within the species range. Population-level differences in egg development, hatching time, and larvae thermal performance in response to temperature (Koeller et al. 2009; Ouellet et al. 2017) suggest local adaptation and thus differential responses to warming temperature.

Along North America, *P. borealis* is primarily distributed on the continental shelf from the Gulf of Maine in the Northwest Atlantic to the Arctic Ocean (Bergström 2000). Within this range, Jorde et al. (2015) identified three distinct genetic groups using ten microsatellites, namely the Gulf of Maine, the Flemish Cap, and the continental shelf spanning the Scotian Shelf to West Greenland. Within this region, bottom temperature appears to be the main driver of genetic population structure with the apparent genetic homogeneity observed at finer spatial scales hypothesized to be a consequence of the geographic scale of larval dispersal (Jorde et al. 2015). Indeed, biophysical modelling studies have also suggested significant geographic connectivity, whereby northern regions (e.g., Arctic and Labrador) supply larval settlers to southern regions (e.g., Newfoundland and Gulf of St. Lawrence) transported by the dominant Labrador Current along the Canadian coast (Le Corre et al. 2019, 2020).

Southern stocks of *P. borealis* are experiencing rapid declines that coincide with warming ocean. The Gulf of Maine stock, for example, experienced significant declines in 2012 associated with an extreme marine heatwave event and predator shift (Whitmore et al. 2013; Richards and Hunter 2021). In Canada, some southern stocks are classified in the critical or cautious zone of their respective precautionary approach frameworks (DFO 2023a, 2023b). The total biomass estimated for the Eastern Scotian Shelf (SFA-13 to 15, Fig. 1) decreased by 26 % between 2021 and 2022 and the stock is considered in the Cautious Zone (DFO 2023b). Similarly, the adult abundance in the Gulf of St. Lawrence (SFA-8 to 12, Fig. 1) showed a downward trend since the mid 2000s (DFO 2023a). In comparison, estimated total biomass of northern most Canadian stocks (i.e., WAZ, EAZ, Fig. 1) was stable over the same period (DFO 2021). Ongoing climate change is expected to further reduce the southern distribution of northern shrimp through significant alterations in habitat, connectivity, and fitness (Le Corre et al. 2021; Leung et al. 2023). Yet, the impact of genomic and adaptive variation across this range, coupled with changing environmental conditions, on this commercially important species remains unclear.

**Figure 1.**
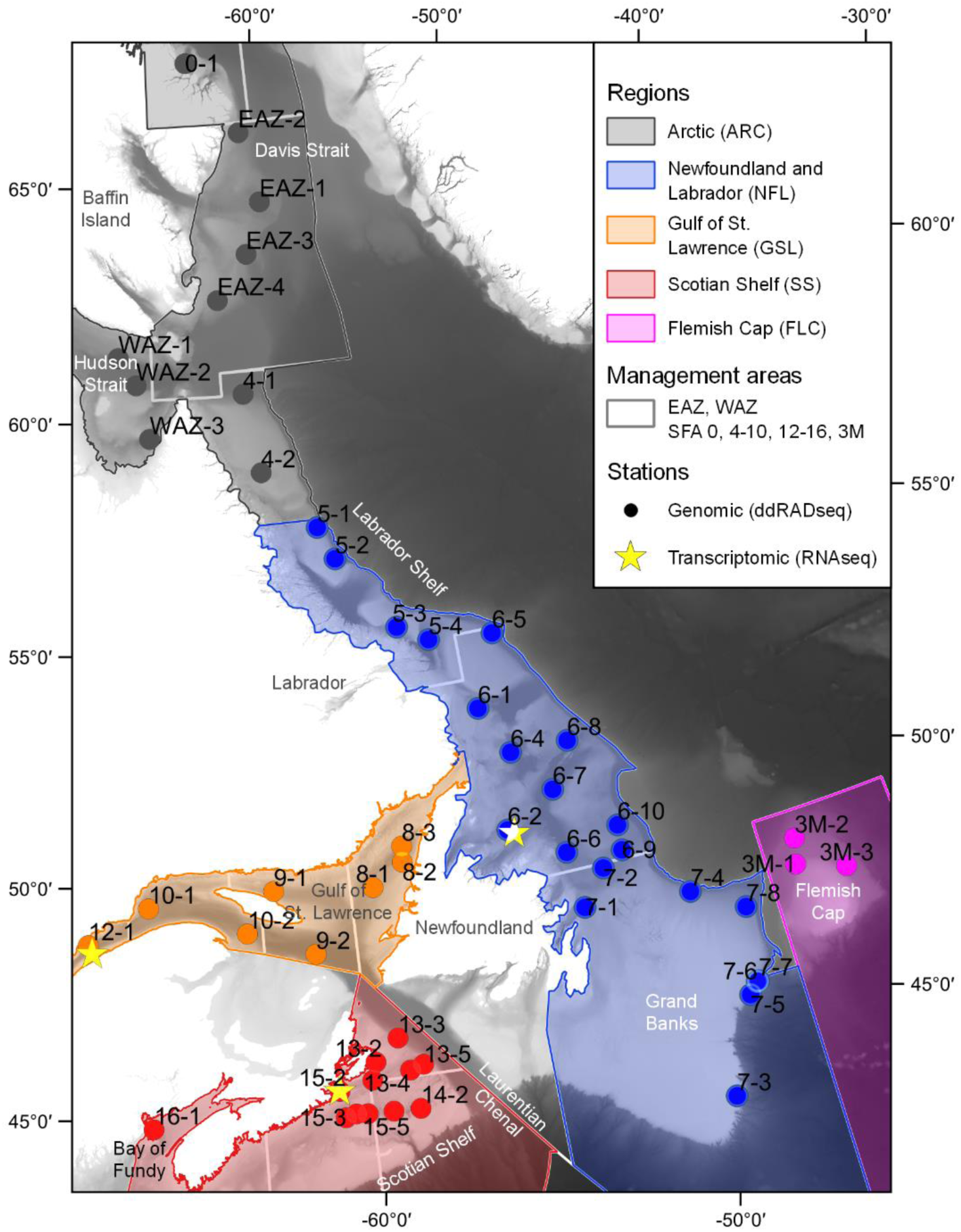
Study area of *Pandalus borealis* from the Bay of Fundy to the Davis Strait, covering five geographic regions and 16 management areas. Stations sampled for the genomic (N = 54) and the transcriptomic (N = 3) datasets are identified with the name of the management areas from Fisheries and Oceans Canada (DFO), i.e., Shrimp Fishing Areas (SFA) 0, 4 to 10, 12 to 16, Eastern Assessment Zone (EAZ) and Western Assessment Zone (WAZ), or Northwest Atlantic Fisheries Organization (NAFO), i.e., 3M (see Table 1 for more details).

**Table 1.** Sampling metadata for the genomic (ddRADseq) dataset. The number (N) of stations and the total N of shrimp sequenced and genotyped (i.e., following removal of misidentified and of low sequencing quality samples) within regions and management areas are presented. All stations were sampled between May and December 2019.

| Region | Management area | N station | N shrimp |  | Sampling month |
| --- | --- | --- | --- | --- | --- |
|  |  |  | sequenced | genotyped |  |
| Arctic (ARC) | SFA-0 | 1 | 14 | 2 | Sept. |
|  | EAZ | 4 | 124 | 108 | Aug. - Sept. |
|  | WAZ | 3 | 71 | 61 | Aug. |
| Newfoundland and Labrador (NFL) | SFA-4 | 2 | 67 | 45 | July - Aug. |
|  | SFA-5 | 4 | 128 | 99 | Oct. |
|  | SFA-6 | 9 | 290 | 254 | Oct. - Dec. |
|  | SFA-7 | 8 | 267 | 254 | May - Nov. |
| Gulf of St. Lawrence (GSL) | SFA-8 | 3 | 116 | 97 | Aug. |
|  | SFA-9 | 2 | 64 | 51 | Aug. |
|  | SFA-10 | 2 | 64 | 57 | Aug. - Sept. |
|  | SFA-12 | 1 | 32 | 26 | Aug. |
| Scotian Shelf (SS) | SFA-13 | 5 | 153 | 150 | June - Sept. |
|  | SFA-14 | 2 | 67 | 66 | June |
|  | SFA-15 | 4 | 120 | 117 | June - Aug. |
|  | SFA-16 | 1 | 16 | 12 | Sept. |
| Flemish Cap (FLC) | NAFO-3M | 3 | 117 | 114 | July |

In this study, we applied high-resolution genomics in the northern shrimp to evaluate factors affecting population structure and leverage this information to predict vulnerability to climate change along the Canadian east coast. To achieve these objectives we 1) characterised population structure using a reduced-representation of the genome approach, 2) inferred spatial and environmental drivers of population structure, 3) identified candidate adaptive loci using genotype-environmental associations and determined their role in temperature response using transcriptomics, and 4) predicted the northern shrimp genomic offset using climate forecasts and assessed its adaptive standing genetic variation (see Fig. S1 for a detailed plan of the objectives, associated datasets, and analyses). Our results provide a novel perspective for how genetic variation and environmental change will influence a species’ response to climate change, offering valuable insights for similarly broadly distributed species.

## Methods

### Sample and environmental data collection

Samples of northern shrimp were collected between May and December 2019 at 54 stations distributed across 16 management areas along the eastern coast of North America (between 2 to 52 shrimps per station, mean ± sd = 31.8 ± 8.9 shrimps per station; Table 1, Fig. 1) and preserved in 95% ethanol. Sample sizes were small at some stations mostly due to the species rarity or misidentification. Stations spanned five geographic regions, i.e., Arctic (ARC), Newfoundland and Labrador (NFL), Gulf of St. Lawrence (GSL), Scotian Shelf (SS) and Flemish Cap (FLC; Table 1, Fig. 1).

Monthly bottom and surface temperature and salinity were extracted for the study area from the Bedford Institute of Oceanography North Atlantic Model (BNAM; Wang et al. 2018). A baseline or ‘current’ climatology was derived as an average of 1990-2015 runs (Wang et al. 2018) and a projected ‘future’ climatology was derived from BNAM simulations based on a high-emissions representative concentration pathway (RCP8.5), for the 2066-2085 period (Brickman et al. 2016). From these climatologies, we derived five averaged environmental variables linked to shrimp life-history (Shumway et al. 1985): 1) winter bottom temperature (BT-w, January to March), 2) summer bottom temperature (BT-s, July to September), 3) surface temperature during larval developmental phase (SST-l, April to July), 4) annual bottom salinity (BS-a), and 5) annual surface salinity (SSS-a). Current environmental values were extracted for each station and were checked for collinearity, which revealed all pairwise correlations <|0.7| (Fig. S2). Variables were centered and scaled (zero mean, unit variance; Table S1) prior analyses.

### Genomic data acquisition

#### Reference genome

One *P. borealis* female was collected in June 2019 within the GSL and flash-frozen in liquid nitrogen. DNA was extracted and nuclear high molecular weight DNA was isolated by BIO S&T (Montreal, Canada). Multiple libraries were prepared and sequenced at Génome Québec (Montréal, Canada) to obtain a variety of read lengths and quality. In brief, two libraries of PacBio sheared gDNA large insert were prepared and sequenced on 16 cells PacBio Sequel sequencing SMRT cell. A 10X Chromium linked-reads library and a shotgun library were prepared and each sequenced on three lanes of Illumina HiSeqX PE150. To improve the first draft assembly obtained (not presented), a second sequencing round was performed for a PacBio HiFi library on a PacBio Sequel II SMRT Cell. The *P. borealis* draft genome was assembled by C3G (McGill University, Montréal, Canada). PacBio sequencing reads were assembled *de novo* using the long-read wtdbg2 (Ruan and Li 2020) and Flye (Kolmogorov et al. 2019) assemblers. Scaffolding was performed using the 10X Chromium linked-reads with ARCS (Yeo et al. 2018) and LINKS (Warren et al. 2015) tools. Finally, the draft genome was polished with the shotgun reads using PILON (Walker 2014). A BUSCO analysis using the lineage dataset arthropod_odb10 (2020-09-10) was performed to assess the quality and completeness of the genome (v.5.4.1, Manni et al. 2021). Genome size was also estimated using the k-mer method implemented in the web version of GenomeScope (Vurture et al. 2017).

#### Single-nucleotide polymorphism (SNP) panels

DNA was extracted with DNeasy Blood and Tissue kits (Qiagen), quality was assessed on a 2% agarose gel, and concentration was quantified on a Synergy LX (BioTek) using PicoGreen. Twenty double digest restriction-site associated DNA (ddRAD) libraries of 96 samples (total 1920 samples) were prepared by the Plateforme d’analyse génomique (IBIS, Université Laval) using 20 ng of DNA per sample and the *PstI* and *MspI* restriction enzymes (see Poland et al. 2012 for more details). Libraries were sequenced on Illumina NovaSeq 6000 S4 PE150 (three flow cells) with 10% PhiX. A total of 42 *P. borealis* samples were replicated as controls, in addition to ten *Dichelopandalus leptocerus*, and ten *Pandalus montagui* added as references for species misidentification (Fig. S3).

Read quality was checked with FastQC (Andrews 2010) and multiQC (Ewels et al. 2016) and adapter sequences were removed with Trimmomatic (v0.39, Bolger et al. 2014). Demultiplexing and quality filtering were performed with the *process_radtags* module of Stacks (v2.4, Catchen et al. 2013; Rochette et al. 2019), with a truncation at 135 bp and a check for the *pstI* restriction site. Reads were aligned to the reference genome using BWA-MEM (Li and Durbin 2009) with default parameters, and coordinate-sorted using Samtools (Danecek et al. 2021). Aligned paired-end reads were assembled with the *gstacks* module and samples with mean coverage < 5x were discarded.

SNPs were first filtered with the *populations* module of Stacks, to only include those present in ≥75% shrimps per management area and with a minor allele frequency (MAF) ≥ 5% for the complete dataset. Then, individuals with > 25% missing SNPs and SNPs with > 10% missing individuals, were removed using VCFtools (Danecek et al. 2021). SNPs not in Hardy-Weinberg equilibrium in ≥ 14 management areas (hw.test function of pegas R package; Paradis 2010; R Core Team 2022) and with median coverage above the 99% quantile (i.e., 25x, computed with VCFr; Knaus and Grünwald 2017) were removed. Linkage disequilibrium (LD) between SNPs was computed intra and inter scaffolds with PLINK (v1.9, Purcell et al. 2007), and one SNP per LD group (r^2^ > 0.5) was selected using a modified version of the radiator *filter.ld* function (Gosselin et al. 2020). Finally, only the first SNP per locus was kept. The final product from the SNP filtration was the “complete SNP panel”. Relatedness coefficients were computed with VCFtools, using the algorithm of Manichaikul et al. (2010), and no related individuals (≥ 0.25) were detected.

SNPs from the complete panel were classified as putative neutral or outlier SNPs using two genome-scan approaches, pcadapt (v4.3.3, Privé et al. 2020) and BayeScan (v2.1, Foll and Gaggiotti 2008). The pcadapt approach is individual-based and identifies outlier SNPs combining principal components analysis (PCA) and Mahalanobis distance. Three principal components were retained based on visual inspection of the scree plot (Cattell 1966). BayeScan is a population-based approach relying on *F*_ST_ distribution between groups, defined here as stations and management areas. SNPs with a *q*-value < 0.05 with either approach were identified as outliers. Two additional SNP panels were then created: 1) the “outlier SNP panel”, and 1) the putative “neutral SNP panel” including SNPs not identified as outliers.

#### Transcriptomics

The transcriptomic data were originally presented in Leung et al. (2023). Briefly, the dataset consists in RNA sequencing data, or transcript counts, from the muscle of 54 adult females from three regions (SS, GSL, NFL; Fig. 1) experimentally exposed to three temperatures (2, 6, 10°C). Differentially expressed transcripts (DETs) were identified for temperature or regional effects (Leung et al. 2023). SNPs associated with transcripts were then identified by mapping the reference transcriptome to the reference genome using Hisat2 (v2.2.1, Kim et al. 2015). Functional annotations of DETs were extracted from the *P. borealis* reference transcriptome (Leung et al. 2023). Enriched gene ontology terms were identified using the Fisher’s exact test implemented in topGO (v2.48, Alexa and Rahnenfuhrer 2022).

### Description of population structure

Genotypic information at the individual level was used to perform PCAs over the three SNP panels with the *glPca* function of adegenet (Jombart 2008). Alleles were identified as 0, 1, and 2 (representing the count of the minor allele) and centered. Missing values were replaced by the mean genotype across all individuals. Minor allele frequencies at the station level were used to compute Euclidian distances and then to perform PCAs using vegan R package (v2.6.2, Oksanen et al. 2022). We also used the Bayesian clustering program Admixture with default parameters (v1.3.0, Alexander et al. 2009) to infer the presence of populations. The optimal number of clusters (K) was obtained with the cross-validation procedure, testing K from 1 to 20.

Indices of genetic differentiation (*F*_ST_) were computed among regions, management areas, and stations using the dartR package (Gruber et al. 2018). Confidence intervals (C.I.s 95%) for *F*_ST_ were calculated using 999 bootstraps. Expected heterozygosity (*He*) was computed per station using the *basic.stats* function of the hierfstat R package (Goudet and Jombart 2022) and differences between regions were assessed with one-way ANOVA’s and Tukey’s HSD tests for multiple comparisons.

### Drivers of population structure

We used Estimated Effective Migration Surfaces (EEMS; Petkova et al. 2016) to identify and visualize variation in gene flow, or effective migration rates (m), based on deviation from the null expectation of isolation by distance under a stepping-stone model. Pairwise genetic dissimilarities were computed using the bed2diffs2 R script (Petkova et al. 2016). The EEMS analysis was run for the three SNP panels to contrast spatial patterns of presumptive gene flow (Storfer et al. 2018). Based on preliminary analyses, we selected 800 demes as the maximum density that adequately reflected geographic patterns within and among regions. The EEMS analysis was run with three chains for 1,000,000 burn-in iterations followed by 3,000,000 MCMC iterations. Default parameters for the EEMS analysis were used except for two changed to optimize acceptance proportions (mEffctProposalS2 = 0.3, mrateMuProposalS2 = 0.005).

The impact of geographic and environmental variables on the distribution of genetic variation was investigated using partial redundancy analyses (pRDAs) following Capblancq and Forester (2021). We used three sets of explanatory variables: 1) neutral genetic structure, 2) geographical variables, and 3) environmental variables. The loadings of the first three axes of the PCA using the neutral SNP panel at the station level were used as proxy of neutral genetic structure. Geographical variables were latitude and longitude at each station. Environmental variables were the five previously described (BT-w, BT-s, SST-l, BS-a, SSS-a). The full RDA model included the three sets of explanatory variables, and MAF at stations as the response variable. Then, pRDA models were used to partition the variance into the relative contribution of neutral genetic structure, geographical, or environmental variables while controlling for the other variables.

Analyses of genotype-environment association (GEA) were also used to identify candidate adaptive loci using pRDAs following Capblancq et al. (2018). We first ran a pRDA with the five environmental variables while controlling for the neutral genetic structure. Then, a Mahalanobis distance was computed using loci loadings of the most informative axis based on a visual observation of the scree plot (K = 2). SNPs with a *q*-value < 3.5×10^−6^ (i.e., alpha level of 0.05 / N SNPs) were identified as candidate adaptive loci.

### Adaptive variation and vulnerability to climate change

Candidate adaptive loci were used to compute an adaptive index across the seascape under current and future environmental conditions following Capblancq and Forester (2021). A RDA with adaptive loci, or an enriched RDA, was built using the allele frequency per station as the response variable, and the five environmental variables as explanatory variables. Then, an adaptive index using the first two axes of the RDA with adaptive loci was computed for each landscape pixel using the loadings and the environmental values.

We also explored genomic vulnerability and adaptive potential of shrimp to changing environmental conditions with two proxies, i.e., the genomic offset and the adaptive standing genetic variation (aSGV). The genomic offset sums the difference between the adaptive indexes under current and future environmental conditions (Capblancq et al. 2020a; Capblancq and Forester 2021). This analysis is spatially explicit and calculated for each map pixel within the study area following the nominal resolution of the climatic model (~1/12^th^ degree). For each station, we extracted the genomic offset of the closest pixel. We also computed an aSGV per station as the allele frequency variance (*p* x *q*) averaged across adaptive loci (Chhatre et al. 2019; Capblancq et al. 2020b).

## Results

### Genomic data for *P. borealis*

The *Pandalus borealis* reference genome assembled has a size of 3.51 Gb, composed of 84,733 contigs with an N50 of 89.5 Kb (Table S2). The *P. borealis* genome size was estimated at 5.03 Gb with the k-mer approach, indicating that the reference assembly is approximately 70% complete (Fig. S4). BUSCO analysis indicated the presence of 67.6% complete orthologs (65.5% single copy, 2.2% duplicated copy), 16.7% of fragmented orthologs, and 15.7% of missing orthologs (n = 1013 orthologs compared). The reference genome is still fragmented despite two rounds of sequencing/assembly with long PacBio reads but of sufficient quality for the mapping of ddRAD reads.

A total of 8.3 billion reads were obtained from all ddRAD libraries with a mean of 4.0 ± 1.9 million reads per sample following demultiplexing and quality filtering (Fig. S1). A catalog of 2.2 million SNPs was created through the *gstacks* module, with a mean effective coverage of 14.6 ± 6.7x. Following filtration steps, the full dataset identified as the complete SNP panel comprised 1,513 individuals and 14,331 SNPs (Table 1; Table S3). The outlier SNP panel was composed of 1,552 SNPs identified with BayeScan (N = 1,188) or pcadapt (N = 1,225) and the majority of outlier SNPs were identified by both approaches (55%). The remaining 12,779 SNPs not identified as outliers by either approach formed the putative neutral SNP panel.

Of the complete SNP panel, 13.4% SNPs (N = 1,916) were associated to at least one transcript (Fig. S5A). This proportion remained relatively stable with 12.8% (N = 198) and 13.4% (N = 1718) for the outlier and the neutral SNP panels, respectively (Fig. S5BC). For SNPs associated to at least one transcript, 43.4%, 43.0%, 46.5% from the complete, outlier, and neutral SNP panels, respectively, were associated with annotated transcripts (Fig. S5A-C). Proportions of 7.1% (N = 135), 7.2% (N = 125), and 5.1% (N = 10) from the complete, outlier, and neutral SNP panels, respectively, were associated to differentially expressed transcript either due to basin temperature or sampling region (Fig. S5A-C).

### Population structure among regions, management areas, and sampling locations

Four genetic clusters were identified using multiple proxies for population structure and the three SNP panels. With the PCA analysis, the FLC region separated from other regions on the first axis at the individual and station levels with all three SNP panels (Fig. 2A-D, S6AB). On the second PCA axis, individuals or stations separated along latitudinal gradient from NFL region to the SS region for both the complete and the outlier SNP panels (Fig. 2BD, S6B). The ARC/NFL regions overlapped partially with the GSL region, which overlapped partially with the SS region using the individual or station level data (Fig. 2BD, S6B). Note that the proportion of variation explained by the first and second PCA axes was an order of magnitude greater using the outlier when compared to the neutral SNP panel at the individual level (6.5% vs. 0.5%; Fig. 2AB).

**Figure 2.**
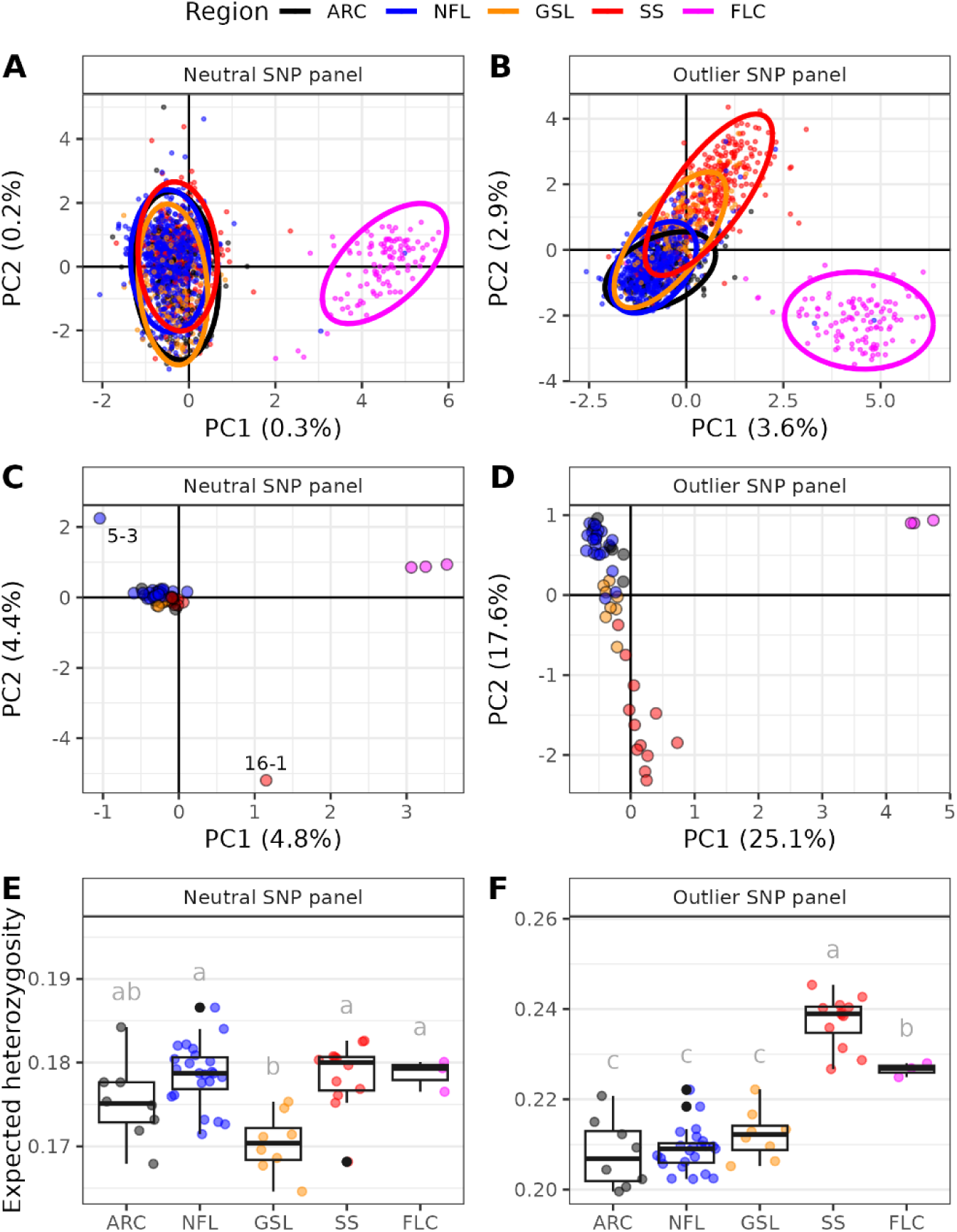
Comparison of population structure of 1,513 *Pandalus borealis* from 54 stations using the putative neutral (A,C,E) and outlier (B,D,F) SNP panels. Principal component analyses (PCA) at the individual (panel A,B) and sampling location levels (panel C,D), and expected heterozygosity computed at the sampling location level (panel E,F). Colour represents the five geographic regions. Expected heterozygosity is presented as boxplots and observed values. Differences in expected heterozygosity between regions were tested using one-way ANOVA (neutral SNP panel: *F*(4,49) = 7.70, *p* < 0.001); outlier SNP panel: *F*(4,49) = 62.53, *p* < 0.001) and Tukey HSD. Stations SFA-0-0, SFA-7-7 and WAZ-3 were excluded from the station level analyses (panel C,D) given their low sample sizes (N < 10). See Fig. S6 for results using the complete SNP panel.

For the Admixture analyses, the cross-validation suggested K=4 for the complete SNP panel, K=3 for the neutral SNP panel, and K=19 for the outlier SNP panel (Fig. S7A). With all SNP panels, the FLC region was distinguished from other regions (Fig. S7B-D). The genetic distinctiveness of ARC/NFL, GSL, and SS regions was less obvious and varied according to the SNP panel used. With the complete and outlier SNP panels, one genetic cluster was more abundant in the ARC/NFL regions, whereas a different genetic cluster was more abundant in the SS region (Fig. S7BD). GSL shrimp had intermediate membership probabilities to the ARC/NFL and the SS clusters with the complete and outlier SNP panels (Fig. S7BD). No more groups were geographically relevant using the outlier SNP panel. The *F*_ST_ estimated between the FLC region and the four other regions were ≥ 0.0198 using the complete SNP panel. Estimates of *F*_ST_ among the four other regions ranged from 0.0006 (ARC vs. NFL) to 0.0036 (ARC vs SS; all *p*-values < 0.001; Fig. S8). Patterns of expected heterozygosity (*He*) differed between SNP panels. In the complete and neutral SNP panels, mean *He* was lower in the GSL region compared to NFL, SS, and FLC regions (*p*-values ≤ 0.02) but was comparable to the ARC region (*p*-values ≥ 0.094; Fig. 2E, S6C). For the outlier SNP panel, the SS and FLC regions had higher mean *He* compared to those of the GSL, NFL and ARC regions (*p*-values ≤ 0.003; Fig. 2F). The SS region also showed a higher mean *He* than that of the FLC region (*p* = 0.003; Fig. 2F).

Finer-scale population structure within regions was investigated at the management area and station levels using indices of genetic differentiation (*F*_ST_), which can be missed using PCA or Admixture analyses. We only presented the complete SNP panel to estimate *F*_ST_ as it provided an averaged signal from neutral and outlier SNPs. Within the ARC and NFL regions, *F*_ST_ ranged from <0.0001 to 0.0012 between management areas, with only two pairwise *F*_ST_ comparisons being not significantly different (SFA 6 and 7, and SFA 5 and 6; Fig. S9). Within the ARC and NFL regions, heterogeneity was also observed between stations as most *F*_ST_ comparisons were significantly different than 0 (Fig. S10BC). Within the GSL region, heterogeneity was observed between management areas; SFA-8 was genetically distinct from other management areas (SFA-10 and 12, *F*_ST_ ≥ 0.0003, *p*-values ≤0.0002; Fig. S9). In fact, stations SFA-8-1 and 2, and not SFA-8-3, were genetically differentiated from most other stations within the GSL region (Fig. S10D). Within the SS region, the four management areas were all distinct from each other (*F*_ST_ range 0.0003 – 0.0023; Fig. S9). The Bay of Fundy (SFA-16) showed the highest *F*_ST_ values of the SS region, both at the management area (all *F*_ST_ = 0.0023; Fig. S9) and station levels (mean *F*_ST_ = 0.0028 ± 0.0006 compared to 0.0008 ± 0.0008 for *F*_ST_ contrasts between other SS stations, *t*(18.2) = 9.74, *p* < 0.001; Fig. S10A). Furthermore, stations SFA-13-3 and SFA13-5 were more genetically similar to stations in the GSL region than those in the SS region (mean *F*_ST_ with GSL stations = 0.0004 ± 0.0002 compared to 0.0015 ± 0.0009 for *F*_ST_ contrasts with other SS stations, *t*(24.9) = −4.66, *p* < 0.001). Shrimp from the SFA-13-3 and SFA13-5 stations were within the GSL cluster in the PCA analysis using the outlier SNP panel (Fig. 2BD). For the FLC region, no genetic differentiation was observed between the three stations (Fig. S10E).

### Drivers of population structure

The EEMS analyses highlighted zones of high connectivity/gene flow or effective migration using the three SNP panels (Fig. 3, S11). Some spatial patterns were similar with the complete, neutral and outlier SNP panels such as the reduced effective migration (i.e., putative barriers to gene flow) detected between the Grand Banks (NFL region) and the FLC region, and within the eastern SS region close to the Laurentian Channel (Fig. 3, S11). Higher effective migration was also detected using all panels within the Hudson Strait (ARC), within the eastern Grand Banks (NLF), within the FLC region, within the western GSL region, and at 45^th^ parallel within the SS region (Fig. 3, S11). Reduced effective migration close to southern Baffin Island (ARC) and increased effective migration along the Labrador Shelf (ARC/NFL) were only observed using the neutral SNP panel (Fig. 3A). In contrast, only the outlier SNP panel suggested a heterogenous pattern of higher and lower effective migration on the southern Labrador Shelf and eastern shelf within the NFL region and lower effective migration in a small coastal area at 45^th^ parallel in the SS region (Fig. 3B).

**Figure 3.**
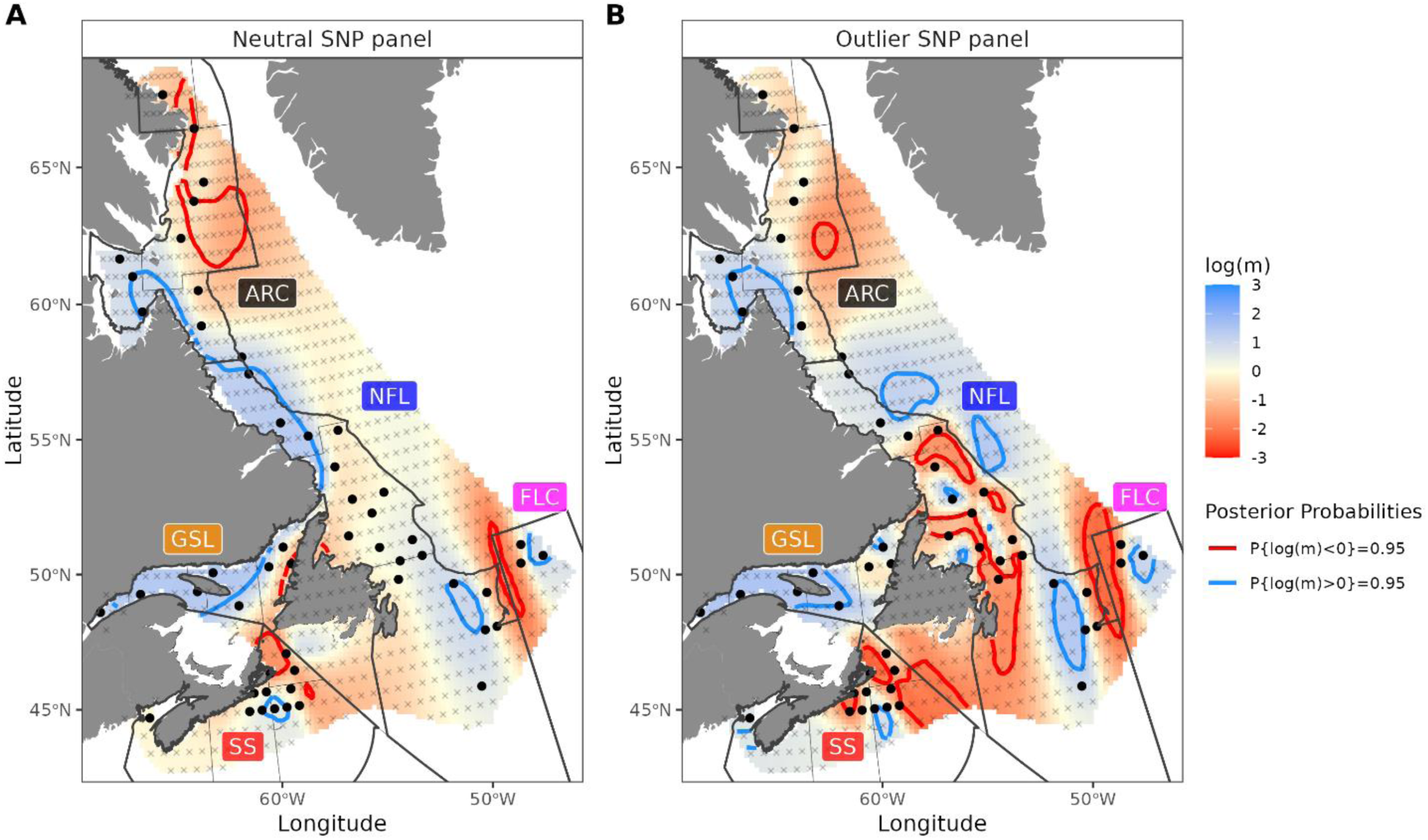
Estimated effective migration (m) surface using the A) neutral and B) outlier SNP panels, combining the results from three independent MCMC iterations. Red areas represent zones of restricted effective migration (i.e., barriers), while blue areas represent zones where migration is higher than expected (i.e., facilitated geneflow). Zones of deviations from the null expectation, i.e., posterior probability P(m ≠ 0 | D) > 95%, are delimited by plain color lines. The gray crosses represent the spatial distribution of demes used (N = 800), while black dots show the stations on deme distribution. Note that the Newfoundland landmass could not be removed from the analysis and a zone of reduced migration was observed with both panels. Results using the complete SNP panel are presented on Fig. S11.

We observed an important proportion of genetic variation explained by neutral genetic structure, environment, and geography with partial redundancy analyses (pRDAs). The full pRDA model explained 35% of the total genetic variance *(p* = 0.001; Table 2). Neutral genetic structure and environmental conditions models explained 12% and 9%, respectively, of the total genetic variance (35% and 26% of explainable variance, *p* ≤ 0.027; Table 2). The geographical model, by itself, explained only 4% of the total genetic variance and did not reach statistical significance (10% of explainable variance; Table 2). Ten percent of the total genetic variance was explained by the three confounded set of variables (28% of explainable variance; Table 2).

**Table 2.**
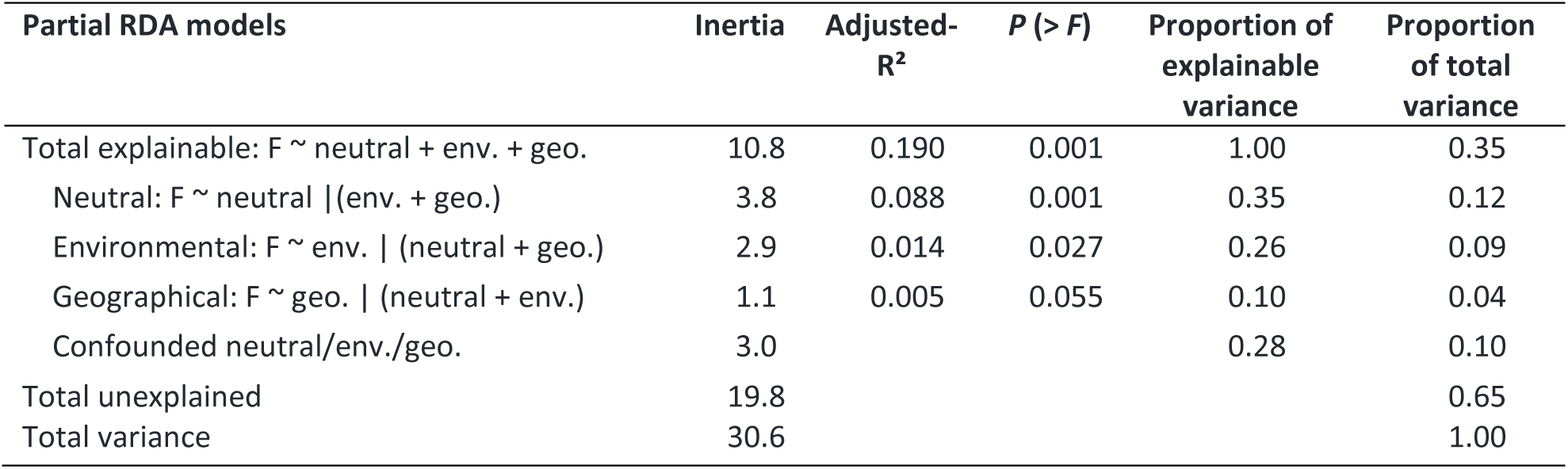
Partial redundancy analyses (pRDA) results partitioning the effect of three sets of explanatory variables (i.e., neutral genetic structure, geographical, environmental). The proportion of explainable variance is the total constrained variation explained by the full model.

The genotype-environment association (GEA) identified 249 candidate adaptive loci from the complete SNP panel with a large proportion of loci (96%, N = 240) previously identified in the outlier SNP panel (Fig. 4AB). Three environmental variables explained a significant portion of the total genetic variance, i.e., bottom or surface salinity during the year (BS-a, 2.6%), surface temperature during larval phase (SST-l, 2.5%) and bottom temperature during winter (BT-w, 1.9%; Table S4). Of the 249 adaptive loci, a proportion of 14.9% (N = 37) was associated to at least one transcript (Fig. S5D). From the latter, around half of the loci (N = 18) were associated to an annotated transcript and only three loci were associated to differentially expressed transcripts (DETs) due to temperature but none of these were annotated (Fig. S5D). A GO enrichment analysis of adaptive loci with annotated transcripts (N = 18) indicated that biological processes linked to signaling between cells or components and response to stimulus were associated to these loci (Fig. S12).

### Adaptive variation and vulnerability to climate change

An enriched RDA on the 249 candidate adaptive loci was conducted to explore seascape adaptive genetic variation under current and future environmental conditions. The adaptive enriched RDA conducted had an adjusted-R^2^ of 0.401 and the first two axes significantly explained the majority of adaptive genetic variance (RDA1: 84.9%, *p* < 0.001; RDA2: 9.2% *p* = 0.048; Fig. 4C). The first RDA axis was related to the surface temperature during larval phase (SST-l), the bottom temperature during summer (BT-s), and bottom or surface salinity during the year (BS-a, SSS-a; Fig. 4C). The second axis was related to all these four variables as well as bottom temperature during winter (BT-w; Fig. 4C). Under current environmental conditions, the first RDA axis revealed a large-scale latitudinal gradient, whereas the second RDA axis incorporated environmental differences mostly associated with depth (Fig. 4D). Adaptive landscapes projected using the 2066-2085 period differed from those of current conditions, with values outside the range obtained with current conditions for the southernmost region (Fig. S13).

**Figure 4.**
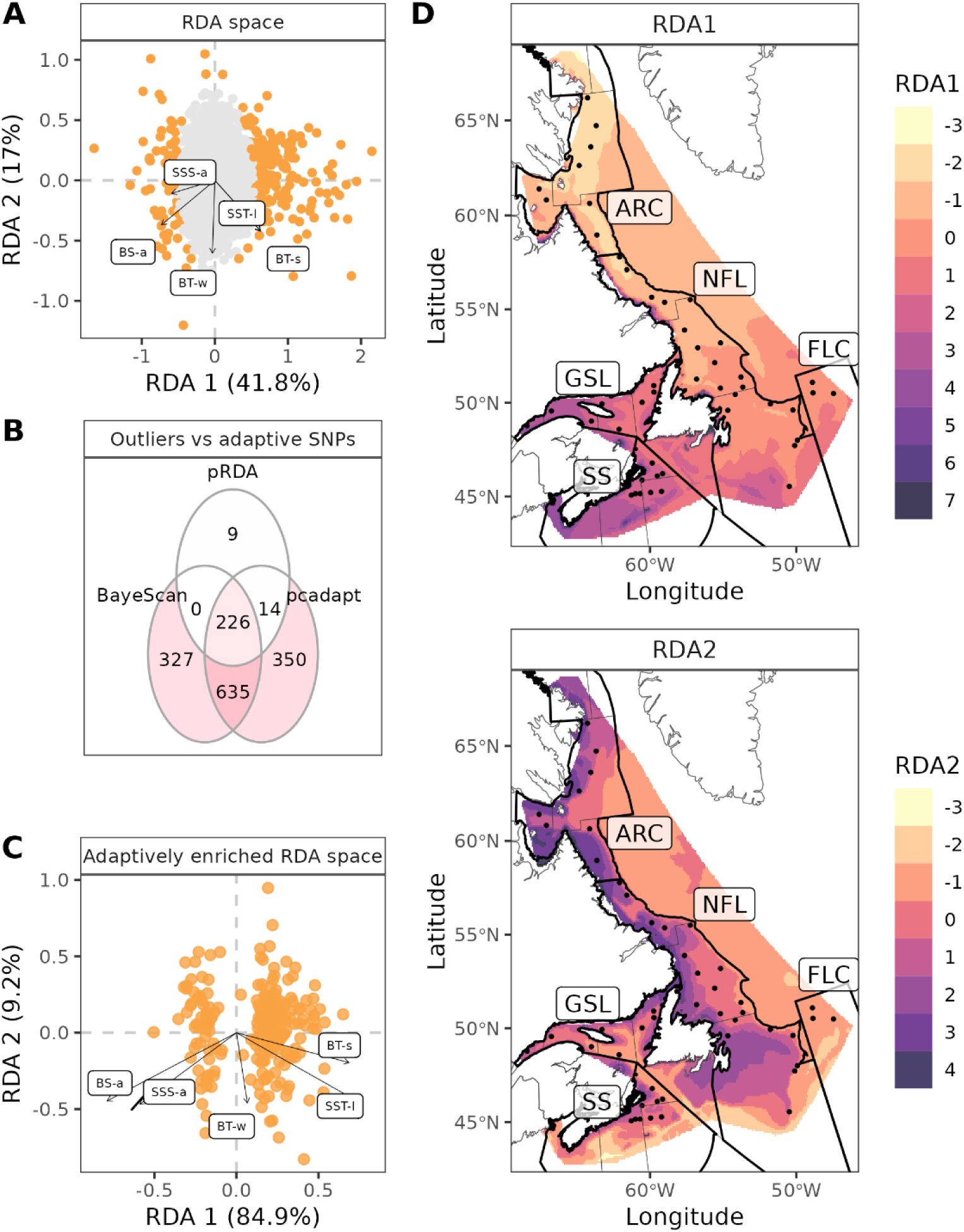
Identification and characterization of candidate adaptative loci (panels A, B) and spatial extrapolation into adaptative landscape under present environmental conditions (panels C, D) in *Pandalus borealis*. A) Projection of SNPs (dots) and environmental variables (vectors) along the first two axes from the partial redundancy analysis (pRDA) constrained using neutral genetic structure. The candidate adaptive SNPs (N = 249) are in orange. B) Venn diagram of SNPs identified as outliers with BayeScan or pcadapt analyses and the candidate adaptive SNPs identified with the pRDA. C) Projection of candidate adaptive SNPs and environmental variables in the adaptively enriched genetic space. D) Spatial projection of the adaptive index into the study area using RDA1 and RDA2. Black dots represent stations, bold lines delimit regions, and thin lines delimit management areas. Close values of adaptive index (e.g., −2 and −1) represent a similar genomic composition associated to environmental variables.

The genomic offset predicted for the 2066-2085 period varies in intensity and variability across regions (Fig. 5A). The genomic offset was generally higher in the southern regions (SS and GSL, mean offset > 0.803) compared to the northern regions (NFL and ARC, mean offset <0.437; Fig. 5A). Moreover, while a spatial gradient in genomic offset was present in the southern regions, the NFL region was relatively homogenous between 50 and 58^°^ latitude (Fig. 5A). The highest genomic offsets were predicted for inshore stations of the eastern SS region (SFA-13-2 and SFA-15-2: Offset ≥ 1.137) and in the Bay of Fundy (SFA-16-1: Offset = 0.993). The lowest genomic offset was predicted for stations in the EAZ management area of the ARC region (EAZ-1: Offset = 0.034). The aSGV also varied between regions (*F*(4,49) = 57.4, *p* < 0.001; Fig. 5B). The SS region had the highest aSVG (mean = 0.169 ± 0.010, *p*-values < 0.001) while the ARC, NFL, and FLC regions showed lowest aSVG (mean range = 0.126 - 0.130, *p*-values > 0.81; Fig. 5B). The genomic offset and aSGV at the station level were positively correlated (*r* = 0.77, *p* < 0.001; Fig. 5C).

**Figure 5.**
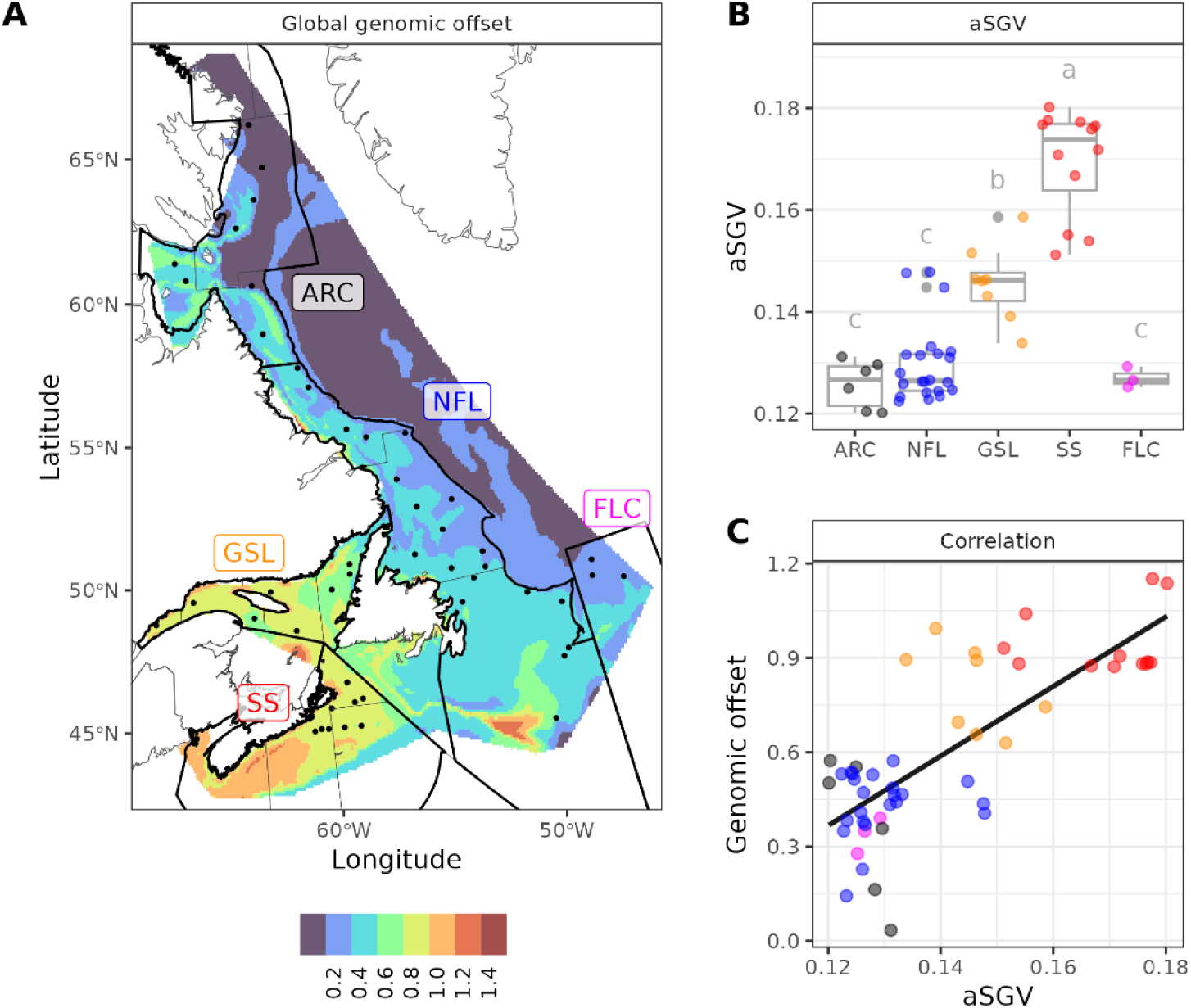
Exploration of *Pandalus borealis* vulnerability to climate change using A) the genomic offset, B) the adaptive standing genetic variation (aSGV), and C) their relationship. A) Genomic offset combining weighted adaptative landscapes of the first two RDA axes under present and future environmental conditions (2075 horizon). B) aSGV computed at station level (dot) and summarized as boxplots. Difference in aSGV between regions was tested with a one-way ANOVA and Tukey HSD. C) Relationship between aSVG and genomic offset extracted for each station (dot, colored by region).

## Discussion

Our study applied high-resolution genotyping in a broadly distributed invertebrate species, revealing complex population structure across multiple geographic scales, explained by broad-scale patterns of connectivity and environmental gradients. We demonstrated that genetic variation and anticipated changes in oceanic conditions led to varying levels of vulnerability, whereby *P. borealis* from southern regions were the most vulnerable to climate change as indicted by the highest genomic offset levels. However, shrimp from the northern regions exhibited less adaptive standing genetic variation (aSVG), suggesting limited evolutionary potential to respond to changing conditions in comparison to other parts of the Northwest Atlantic. Together our results provide a novel insight into the joint role of connectivity and adaptation in the current distribution of *P. borealis* and a first perspective of climatic vulnerability in boreo-Arctic meroplanktonic species.

### Neutral and outlier SNPs uncover population structure at multiple scales for *P. borealis*

Our results identified four large genomic clusters, or populations, in the Northwest Atlantic and Eastern Arctic, including the Flemish Cap (FLC), the Arctic to Newfoundland Shelf (ARC/NFL), the Gulf of St. Lawrence extending to the Laurentian Channel (GSL), and the Scotian Shelf (SS). These results support those from Jorde et al. (2015) using microsatellites for the presence of a population on Flemish Cap, but also reveal genetic differentiation and complexity along the continental shelf using thousands of genome-wide SNPs. Multiple studies exposed structure using population genomics in marine vertebrates or invertebrates in the Northwest Atlantic, challenging the paradigm of general panmixia for many marine species (e.g., Benestan et al. 2015; Van Wyngaarden et al. 2017; Kess et al. 2021; Dorant et al. 2022; Ferchaud et al. 2022; Fuentes-Pardo et al. 2023; Jones et al. 2023). We build on this body of evidence with an uniquely high genomic- and geographic-resolution study for a meroplanktonic species sampled across their northwest Atlantic distribution in addition to samples in the Eastern Arctic. The presence of four genomic populations in our study area despite the large dispersal potential of northern shrimp highlights that other species with similar distributions and plantonik life histories history strategies (e.g., *Pandalus montagui*, *Hyas araneus, H. coarctatus*, *Lebbeus groenlandicus, Eualus gaimardii, E. belcheri*) can also present unexpected broad-scale patterns of population structure. The unprecedented level of genomic information from this study is highly valuable for these species that are less abundant or of lower economic importance.

The large-scale population structure for *P. borealis* reflects its post-glacial recolonization along North America. We showed three *P. borealis* populations between the 45^th^ and the 50^th^ parallels and only one population north of the 50^th^ parallel. This pattern is in line with genetic diversity generally observed in boreal regions where species expanded their range from their southern distribution following the Last Glacial Maximum (Shaw et al. 2002). The poleward colonization occurred faster in the southern regions compared to the northern regions, giving time for drift and divergence to occur in southern regions and more populations to form. We also observed lower heterozygosity at neutral loci in the GSL region (Fig. 2E). Reduced heterozygosity is generally due to a founder effect or a bottleneck following a range expansion leading to lower diversity across the genome (Hewitt 1996, 2000). Other species from the GSL area, often excluded from large-scale studies (e.g., Bradbury et al. 2010; Jorde et al. 2015; Giakoumis et al. 2023), might also possess a distinct genomic signature of past demographic history.

Our study offers a highly detailed analysis of structure in the Northwest Atlantic, revealing distinctions both between and within existing shrimp management areas. The extensive dataset and cost-effective genotyping-by-sequencing method (ddRAD) were instrumental in achieving this level of granularity. We found weak but extensive structure throughout the region, sometimes differentiating adjacent management areas (Fig. S9). Genetic differentiation was also observed within management areas, even between stations separated by < 35 km (e.g., SFA 13-4 and 13-5, *F*_ST_ = 0.0007, *p* = 0.003). The genetic differentiation was often associated with bathymetric features limiting current between adjacent zones such as the Canso hole, the Misaine hole, and the Louisbourg hole in the eastern SS region (Koeller 2000). Genetic differentiation and reduced effective migration between inshore and offshore locations (e.g., SFA 15 in SS region, SFA6 in NFL region) was also observed in Norwegian populations of *P. borealis* of the Northeast Atlantic (Hansen et al. 2021). These patterns of genetic differentiation detected at small spatial scales are in contrast to the presumptive scale of dispersal potential for this meroplanktonic species (e.g., Le Corre et al. 2019, 2020), pointing to yet the potential for local retention for *P. borealis* and other meroplanktonic species.

Northern shrimp from the southernmost management area in the Bay of Fundy (SFA-16) showed the highest level of genetic differentiation observed between management areas within a region. Jorde et al. (2015) previously described another population in the Gulf of Maine (GOM), located south of our study area using microsatellites. It is highly probable that the shrimp from the Bay of Fundy belong to the GOM group identified by Jorde et al. (2015). The high connectivity between the GOM and the Bay of Fundy has been shown in fish and large invertebrate species such as Atlantic cod (*Gadus morhua*; Puncher et al. 2021a, 2021b), Atlantic haddock (*Melanogrammus aeglefinus*; McCracken 1960; Fowler 2011) and American lobster (*Homarus americanus*; Incze et al. 2010; Quinn et al. 2017). However, most of the GOM population previously identified is currently severely depleted (ASMFC 2021) likely due to warming water temperatures, extreme weather events including a heatwave that occurred in 2012, and a shift in predator distribution (Richards and Hunter 2021). *P. borealis* are also found in very low densities in the western Scotian Shelf likely due to unfavorable temperature (Koeller 2000), with multiple tows were necessary to catch an even low number (N = 12) in the Bay of Fundy for our study.

### Connectivity and environmental variation as drivers of population structure

Neutral population structure explained the largest part of the genetic variation in *P. borealis,* representing patterns of broad connectivity during the larval phase in our study area. Indeed, our findings of facilitated or reduced effective migration (Fig. 3A) at a broad geographic scale were generally in agreement with results from biophysical particle-tracking modelling studies and knowledge on the prevailing surface oceanic currents (Le Corre et al. 2019, 2020). The low connectivity between the Grand Banks and Flemish Cap matches the general ocean circulation persistently going around the Flemish Cap (Stein 2007) that limits larval exchange and serves to isolate the bank from adjacent shelf areas (Le Corre et al. 2020). In contrast, a zone of high connectivity in the EEMS analysis was identified from the Hudson Strait to the Labrador Shelf. In this area, surface circulation is mainly driven by the Labrador Current running north to south along the shelf. Biophysical modelling of larval dispersal in this area showed high connectivity from the southern ARC to the NFL region (Le Corre et al. 2020). Similarly, the genetic differentiation observed between the NFL and the GSL regions or between the NFL and the SS regions correspond with previous modelling studies that suggested limited biophysical exchange between these regions (Quinn et al. 2017; Le Corre et al. 2019, 2020).

Environmental variation was the second most important factor explaining the genetic variance of *P. borealis* across the study area (Table 2). In this case, the relationship between environmental conditions and genetic variation may reflect adaptation to local conditions, which impart heterogeneous patterns of selection. The important effect of environmental gradients on genetic structure was also described in past invertebrate studies (e.g., Jeffery et al. 2018; Mendiola and Ravago-Gotanco 2021; Dorant et al. 2022). Here, the wide latitudinal scale and the high geographic resolution of our study allowed us to disentangle the effect of multiple environmental conditions across a considerably larger geographic extent with greater explanatory power than previous studies in the region. At this scale, the environmental variable explaining the largest proportion of genetic variation in *P. borealis* was mean bottom salinity (BS-a; Table S4), which contrast with previous studies identifying temperature as a main driver of local adaptation (Jorde et al. 2015; Stanley et al. 2018). While the effect of salinity is rarely investigated in *P. borealis* (Chang et al. 2021), it is identified as an important factor underlying the genetic structure in other marine invertebrates (e.g., Atlantic scallop, Lehnert et al. 2019; American lobster, Dorant et al. 2022) and an important driver of local adaptation (Defaveri and Merilä 2014; Wenne et al. 2020). Northern shrimp are encountered across a range of salinity, with a broad-scale gradient caused by oceanic currents, but also on finer scale gradients linked to freshwater supply in estuarian and coastal areas (Fig. S2C), which may favour important selection process.

Surface temperature, and to a lesser extent bottom temperature, also explained genetic variation in northern shrimp (Table S4). For northern shrimp, dispersal primarily occurs during the larval phase and the mean surface temperature during the larvae period (SST-l) had the strongest relationship with genetic variation. This pattern has also been observed in other invertebrate species, including the orange mud crab (*Scylla olivacea*), where surface temperature during spawning season best explained observed genetic variation (Mendiola and Ravago-Gotanco 2021). For many marine species, temperature plays a key role during their larval period with a strong influence on key traits such as pelagic larval duration (Quinn et al. 2013), behaviour (Stanley et al. 2016), and mortality (Quinn 2017), all of which interact with oceanographic currents to influence connectivity potential. Large variation in mean SST-l was observed in our study area, ranging from near 0°C in the Arctic to up to 10°C in the Bay of Fundy (Fig. S2C). Correspondingly, differences in larval survival and growth observed in populations of northern shrimp implied the potential for local adaptation to gradients in surface temperature experienced by larvae in the spring-early summer (Ouellet et al. 2017). Interestingly, SST-l explained 30% more variance than the winter bottom temperature (BT-w). The strong gradient in BT-w across the Scotian Shelf and within the study region has been linked to observed genetic structure in past genetic studies of northern shrimp (Jorde et al. 2015; Stanley et al. 2018), Atlantic scallop (Lehnert et al. 2019), and American sandlance (Jones et al. 2023), suggesting that post-settlement processes are important in adaptive variation. Our results build on this observation by showing that winter bottom temperature was an influential factor in genetic structure, but that selection pressures imposed by gradients in surface temperature experienced during the larval period may be more influential for *P. borealis* in our study region.

The important effect of temperature on gene expression of adult *P. borealis* was confirmed through a common garden experiment (Leung et al. 2023). Here, we identified that 14.9 % (N = 37) of the 249 adaptive loci were associated to at least one transcript, which is a similar proportion for the complete SNP panel (13.4%). Signal transduction was the main biological process associated with adaptive loci transcripts (Fig. S12). Interestingly, the same biological process was also observed in a recent genomic study on the sea cucumber *Apostichopus californicus*, for which sea surface environmental variables were most correlated with genetic variation and small proportion (9%) of candidate adaptive SNPs matched to gene ontologies (Lowell et al. 2023). The latter result is expected as neither *A. californicus* nor *P. borealis* are model species (Leung et al. 2023; Lowell et al. 2023). Nonetheless, we did not observe a difference in proportions of SNPs associated to DETs in adult shrimp between the adaptive, the neutral and the outlier SNP panels. In addition, only three adaptive loci were associated to DETs. These results suggest that the adaptive loci identified in our study may not be associated to functions under selection at the adult stage. This interpretation is consistent with our analyses that indicate pelagic environmental variation was best associated with observed genetic variation, implying that selection during the pelagic larval phase may play a more significant role driving adaptive differentiation than post-settlement processes.

### Uneven vulnerability to climate change in the Northwest Atlantic

Species distribution modelling is a key tool for ecologists seeking to predict the impact of changing environmental conditions on species distribution. While offering a robust statistical framework to assess the influence of environmental variation on distribution, this approach can have reduced accuracy at large spatial scales when species’ response to the environment is assumed to be spatially homogeneous (Lowen et al. 2019). Building evolutionary information into predictions of species response to climate change can be particularly important in the presence of strong environmentally mediated genetic structure. One approach to accomplish this integration is through the application of a genomic offset analysis or a space-for-time substitution model, which provides visualization of genomic vulnerability using adaptive genetic variation projected under current and future environmental conditions (Capblancq et al. 2020a; Rellstab et al. 2021; Layton and Bradbury 2022). Genomic offset analyses conducted for our study area showed that regions and management areas in the southern extent of our study (i.e., SS, GSL) have the largest genomic offset, indicating that they are most vulnerable or least adapted to anticipated future conditions. This result aligns with assessments for shrimp in the southern extent of their range where populations have exhibited declining abundance and appear already negatively affected by climate change (see also Bay et al. 2018). The GOM population is severely depleted (ASMFC 2021), and recent abundance indices in the SS and GSL regions indicate important declines in biomass over the last decade (DFO 2023a, 2023b). Increased temperature was identified as an important factor affecting the survival of pelagic larval stages and adults of *P. borealis* in common garden experiments (Ouellet et al. 2017; Leung et al. 2023; Guscelli et al. 2023). It is important to note that genomic offset analyses do not explicitly account for demographic processes such as gene flow and assume an equilibrium in allele-environmental frequencies (Waldvogel et al. 2020; Rellstab et al. 2021), which may limit their ability to accurately reflect genomic variation in vulnerability to climate change. Agreement between the range-wide genomic offset analyses and controlled common garden experiments provides some independent validation of the space for time substitution approach applied in this study. Assessments of species climatic vulnerability are significantly strengthened with the inclusion of multiple approaches, which collectively provide a more comprehensive assessment of biological responses to changing environmental conditions.

We also assessed the ability of sampled populations to respond to changing conditions and selective pressures by measuring the adaptive standing genetic variation (aSVG). In this context, a population with larger aSVG has greater evolutionary potential or is more likely to be able to adapt to changing environments (Barrett and Schluter 2008; Bernatchez 2016). For northern shrimp, the SS region had the greatest aSGV compared to all other regions, followed by the GSL region. Fluctuating selection due to variable environmental conditions can help maintain genetic diversity (Bell 2010; Bernatchez 2016). The amplitude of fluctuations in environmental conditions is uneven across the Northwest Atlantic, with SS and GSL regions experiencing larger variation in temperature and salinity compared to areas at higher latitudes (Weare and Newell 1977; Petrie 2007). Higher levels of aSGV can also be attributed to gene flow between populations experiencing different selective pressure or environmental conditions (Barrett and Schluter 2008; Bernatchez 2016), though in this case, it would be expected that the GSL region would have a higher aSGV given it is adjacent to SS to the south and NFL to the north. While the mechanism behind the greater aSVG in the southern part of our study region cannot be definitively pinpointed, our results suggest higher evolutionary potential in southern *P. borealis* populations.

The genomic offset and common garden experiments reveal that southern regions of *P. borealis* are particularly susceptible to climate change. Paradoxically, the aSGV indicates that animals from these vulnerable regions also possess the highest evolutionary potential. This juxtaposition creates a form of “double jeopardy” for *P. borealis*, emphasizing unexpected challenges in the face of current and ongoing climate change. Although the southern regions already experience suboptimal environmental conditions, these populations, despite its high adaptive genetic variation, likely lack the metabolic functions needed to withstand warmer conditions (Ouellet et al. 2017; Leung et al. 2023; Guscelli et al. 2023). In contrast, *P. borealis* from NFL and ARC regions are relatively less vulnerable to climate change based on the genomic offset, while at the same time presenting lower adaptive genetic diversity (Fig. 5AB). Given larval dispersal is the predominant vector for dispersal, it is unlikely that gene flow from shrimp with more aSGV or adapted to future conditions will occur towards higher latitudes over short time scales given it is countercurrent to the dominant north to south Labrador Current along the coast (Le Corre et al. 2020). Consequently, our results showing the widespread intermediate genomic offset values and the lower aSGV of the NFL region suggest a vulnerability to projected warming conditions with potential for declining patterns in biomass characteristic of lower latitude regions. Our predictions leveraging genotype-environment associations combined with results from common garden experiments should be further explored to characterize the extent of this impact (Waldvogel et al. 2020). Still, insights from genomics presented and discussed provide important context for the status of this species that should be incorporated into management decisions in the SS, GSL, and NFL regions, and a rare perspective for other invertebrates with similar distributions, with less economic importance but essential to the ecosystem functioning, that may be affected comparably by future environmental conditions.

## Acknowledgement

We are grateful to the Ministerio de ciencia e innovaccion and Instituto Espanol de oceanografia for sampling in Flemish Cap area and to the Northern Shrimp Research Foundation for collection of samples from the Arctic and Northern Labrador Shelf, as well as numerous Fisheries and Oceans Canada staff for collection of samples during their regular surveys. We thank F. Lefebvre and G. Leveque (C3G) for the reference genome advice and assembly, and E. Parent, G. Cortial, J. Larivière, L.-A. Dumoulin, F. Paquin and A. Atikessé for laboratory work and sample coordination (DFO, Canada). We would also like to thank Claude Nozères for helpful discussions on crustacean meroplanktonic species inhabiting similar marine depths as *P. borealis*. This project was funded by Genomics Research and Development Initiative (GRDI) of the Government of Canada.

## Data accessibility

Raw sequence data for the *P. borealis* reference genome assembly, the ddRAD and the transcriptomic datasets will be available in the Sequence Read Archive (SRA) under the BioProject accession PRJNA909194 upon the acceptance of the manuscript. Metadata and scripts to obtain SNP panels from raw reads and perform analyses will be available on Zenodo upon the acceptance of the manuscript.

## Author Contributions

AB, NWJ, RRES and GJP designed the study. AB produced the SNP data. AB analysed the SNP genomic data and CL analysed the transcriptomic data. AB wrote the first draft of the manuscript. All authors discussed and interpreted and results and contributed to the writing.

