## Supplemental material for "Diving into broad-scale and high-resolution population genomics to decipher drivers of structure and climatic vulnerability in a marine invertebrate"

**Table S1** **Description of environmental variables observed in the study area.** Mean bottom temperature during winter (BT-w) and summer (BT-s), sea surface temperature during larval developmental phase (SST-l), annual sea surface salinity (SSS-a) and bottom salinity (BS-a).

| Variable | Range | Mean | SD |
|---|---|---|---|
| **BT-w** | -0.72 – 5.66 | 2.75 | 1.58 |
| **BT-s** | -1.20 – 12.46 | 3.08 | 2.23 |
| **SST-l** | 0.15 – 10.10 | 4.20 | 2.53 |
| **BS-a** | 29.53 – 34.94 | 31.18 | 0.98 |
| **SSS-a** | 19.51 – 33.88 | 33.87 | 2.52 |

**Table S2** **Newly created *Pandalus borealis* reference genome statistics.** All statistics are based on contigs of size >= 3,000 bp, unless otherwise noted.

| **Statistic** | **Draft genome** |
|---|---:|
| # contigs (>=0bp) | 84,733 |
| # contigs (>=1,000bp) | 82,165 |
| # contigs (>=5,000bp) | 66,342 |
| # contigs (>=10,000bp) | 56,556 |
| # contigs (>=25,000bp) | 38,419 |
| # contigs (>=50,000bp) | 22,711 |
| Total length (>=0bp) | 3,509,123,860 |
| Total length (>=1,000bp) | 3,507,330,092 |
| Total length (>=5,000bp) | 3,459,619,483 |
| Total length (>=10,000bp) | 3,387,936,425 |
| Total length (>=25,000bp) | 3,086,522,907 |
| Total length (>=50.000bp) | 2,516,262,261 |
| # contigs | 74,313 |
| Largest contig | 993,695 |
| Total length | 3,491,403,202 |
| GC (%) | 36.98 |
| N50 | 89,516 |
| N75 | 45,603 |
| L50 | 11,273 |
| L75 | 24,853 |
| # N’s per 100 kbp | 1.57 |

**Table S3 Details of filtration steps used to obtain the full SNP panel for *P. borealis* from 54 stations in North America distribution.** 145 individuals from the transcriptomic experiment were removed from the final dataset.

| Filter | Individuals retained | RAD loci retained | SNPs retained |
|---|---|---|---|
| Post-alignment *P. borealis* individuals | 1,841 | | |
| gstack: 5x coverage | 1,722 | 886,087 | |
| population: R = 0.75, r = 0.75, MAF > 0.05 | 1,722 | 24,528 | 56,223 |
| VCFtools: Individual > 25% and SNPs > 10% missing | 1,658 | 17,807 | 36,207 |
| Pegas: HW disequilibrium computed by areas | 1,658 | 17,093 | 34,983 |
| VCFtool: Individuals with relatedness > 0.25 | 1,658 | 17,093 | 34,983 |
| VCFr: SNPs with median coverage > 25 (quantile 99%) | 1,658 | 16,764 | 34,485 |
| Plink: SNPs with LD r2 > 0.5 | 1,658 | 14,667 | 20,136 |
| Final: 1 SNP by RAD loci, selected samples and MAF >=0.05 | 1,513 | 14,331 | 14,331 |

**Table S4 Partial redundancy analysis results partitioning the influence of each of the five environmental variables (i.e., BT-w, BT-s, SST-l, BS-a, SSS-a).**

| Partial RDA models | Inertia | Adjusted-$R^2$ | *P* (> *F*) | Proportion of explainable variance | Proportion of total variance |
|---|---|---|---|---|---|
| Environmental: F ~ env. \| (neutral) | 3.69 | 0.042 | 0.001 | 1.00 | 0.120 |
| Winter bottom temp.: F ~ BT-w \| (others env. + neutral) | 0.58 | 0.003 | 0.030 | 0.16 | 0.019 |
| Summer bottom temp.: F ~ BT-s \| (others env. + neutral) | 0.57 | 0.003 | 0.101 | 0.15 | 0.019 |
| Larvae period surface temperature: F ~ SST-l \| (others env. + neutral) | 0.76 | 0.010 | 0.001 | 0.21 | 0.025 |
| Annual bottom salinity: F ~ BS-a \| (others env. + neutral) | 0.80 | 0.011 | 0.001 | 0.22 | 0.026 |
| Annual surface salinity: F ~ SSS-a \| (others env. + neutral) | 0.52 | 0.001 | 0.401 | 0.14 | 0.017 |
| Confounded | 0.46 | | | 0.13 | 0.015 |
| Total variance | 30.61 | | | | 1.000 |

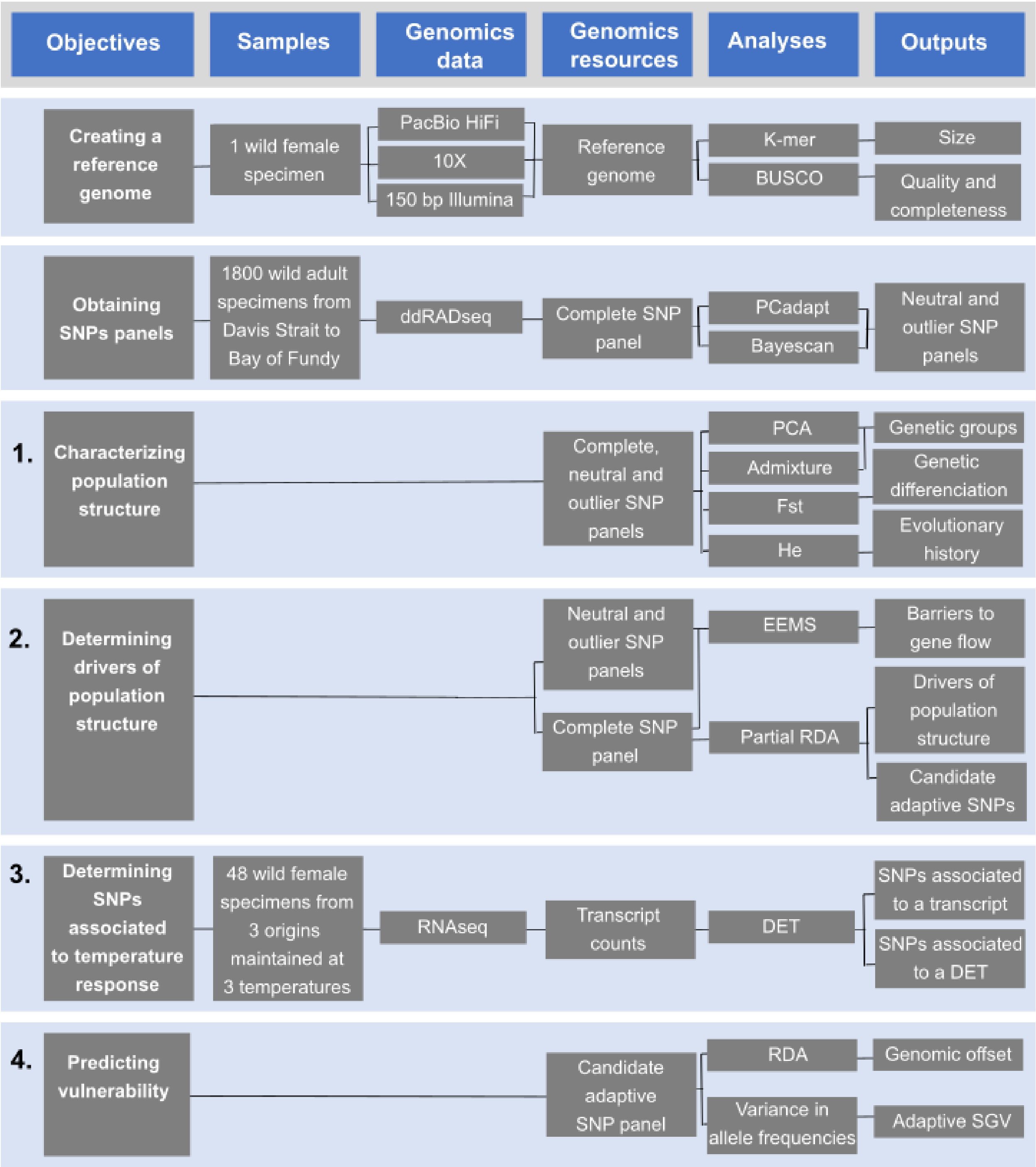


**Figure S1** **Roadmap of the analysis performed in this study to reach the four main objectives.** Biological samples used, genomic data and resources acquired and analysis outputs are presented.

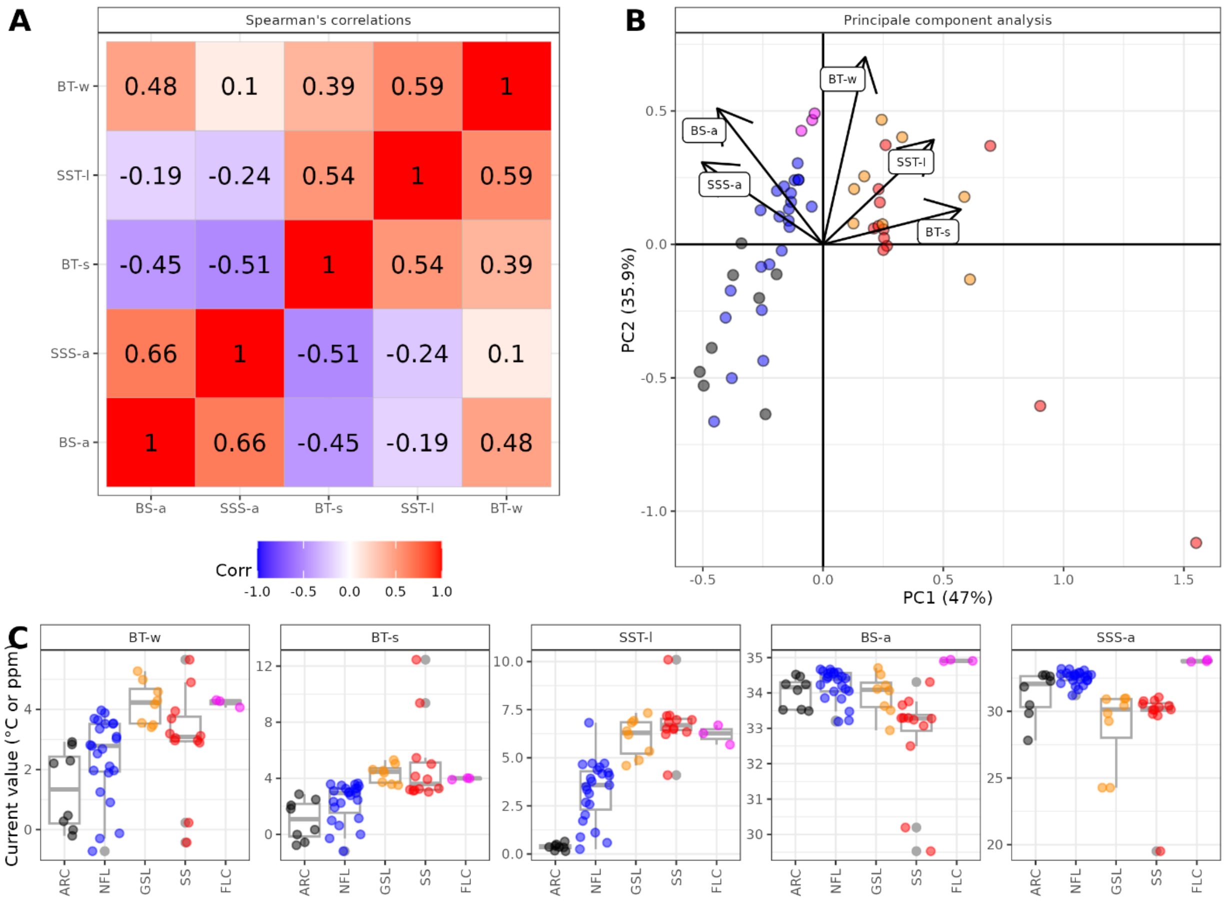


**Figure S2 Relationship among stations and environmental variables. A) Pearson correlation, B) principals component analysis (PCA) and C) distribution of environmental variables for current climatology extracted from BNAM for the 54 stations.** Environmental variables are mean bottom temperature during winter (BT-w) and summer (BT-s), sea surface temperature during larval developmental phase (SST-l), annual sea surface salinity (SSS-a) and bottom salinity (BS-a). Panel C presents both observed values (colored points) and boxplots (Q1,median, Q3 and outliers in gray) by sampling regions. Point color in the panel B and C represents sampling regions (Arctic (ARC): black; Newfoundland and Labrador (NFL): blue; Gulf of St. Lawrence (GSL): orange; Scotian Shelf (SS): orange; Flemish Cap (FLC): pink.

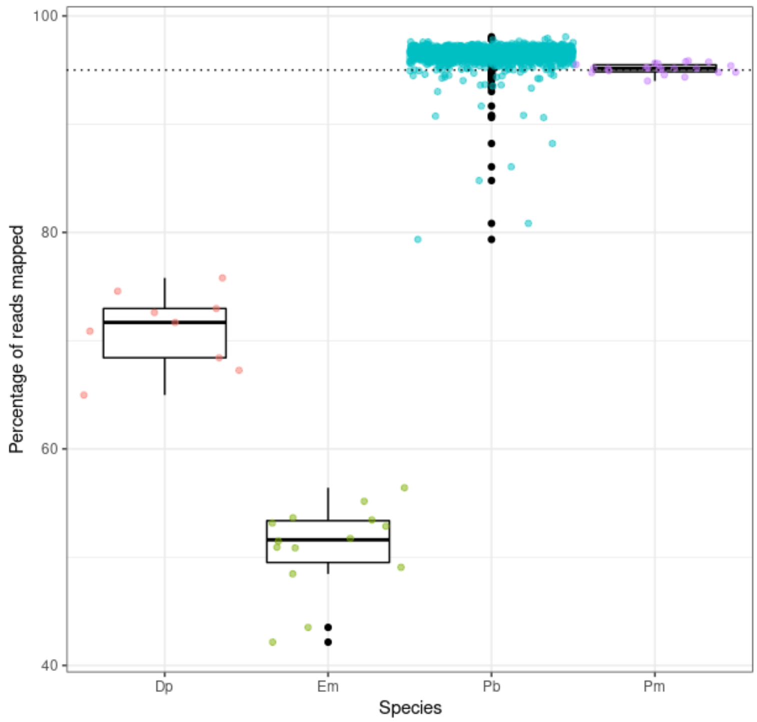


**Figure S3 Percentage of ddRADseq reads mapped to the new *P. borealis* genome observed for different species (Dp: *Dichelopandalus sp.*, Em: *Eualis macilentus*, Pb: *Pandalus borealis* and Pm: *Pandalus montagui*).** Dashed line represents the threshold of 95% of mapped reads to keep a *P. borealis* individual in the analysis. A total of 23 individuals incorrectly identified as *P. borealis* were removed (13 reclassified as *Eualus macilentus* and 11 as *P. montagui*) based on low mapping rate with the reference genome and supplementary COI analysis.

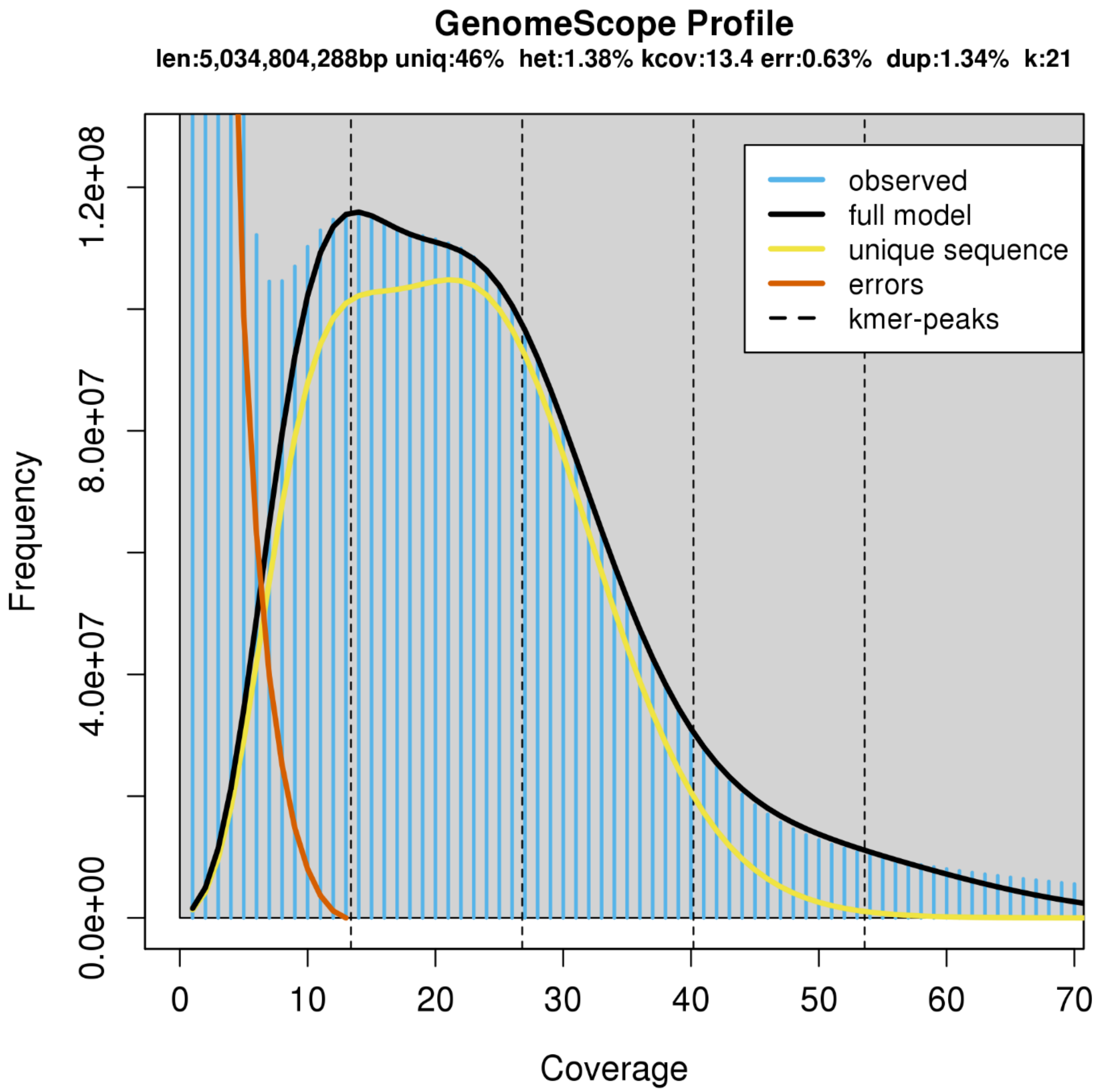


**Figure S4** Genoscope profile obtained from k-mer distribution computed over Illumina short reads with Jellyfish v2.2.3 (using a k-mer length of 21 (Marçais and Kingsford 2011).

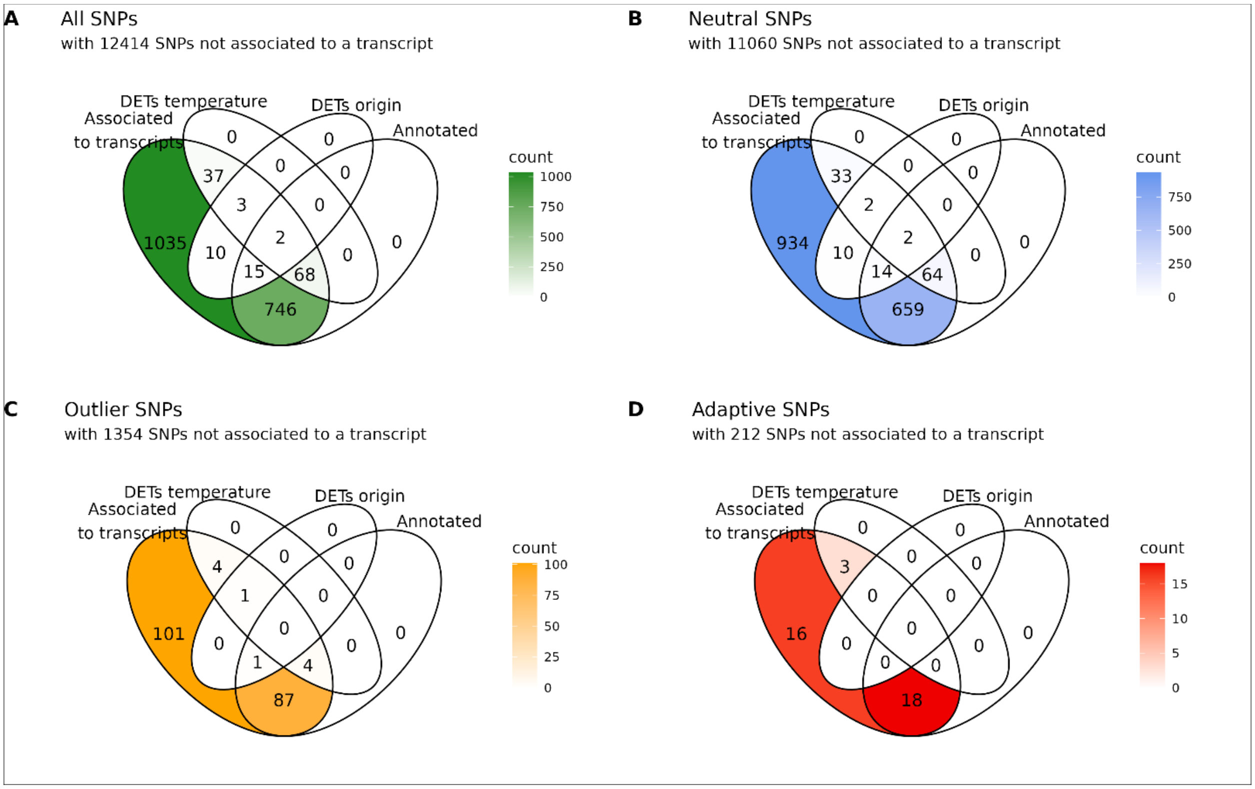


**Figure S5 SNPs associated to a transcript. Venn diagrams representing the number of SNPs associated to a transcript, for the A) complete, B) neutral, C) outlier and D) adaptive SNP panels.** For each panel, the number of transcripts associated to differentially expressed transcripts across laboratory temperature (DETs temperature) and shrimp region (DETs origin), as well as those associated to functionally annotated transcripts are specified.

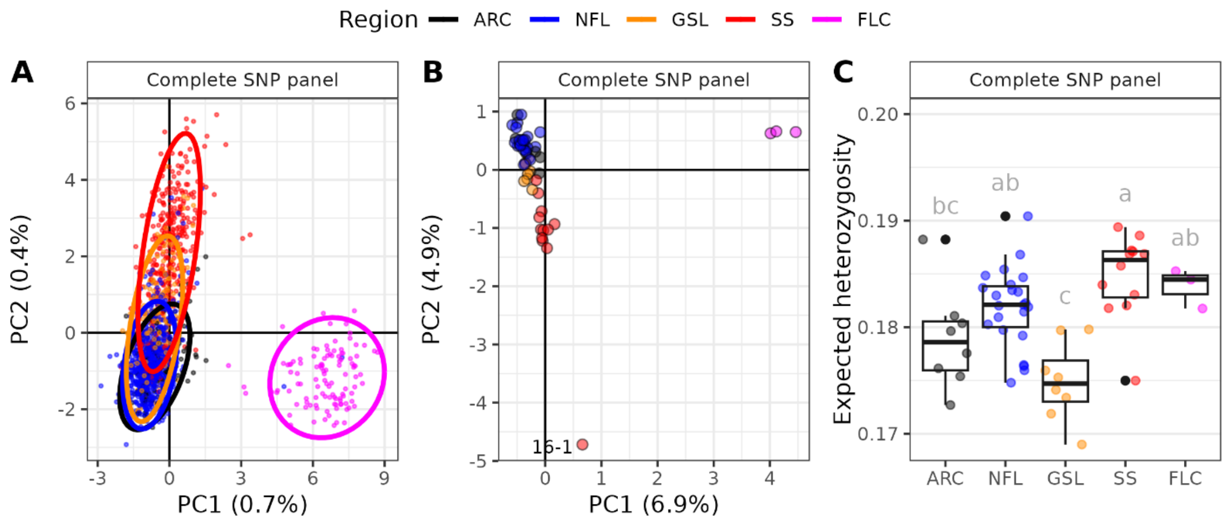


**Figure S6** **Comparison of population structure of 1,513 *P. borealis* individuals from 54 stations using the complete SNP panels.** Principal component analyses (PCA) at the individual (panel A) and station levels (panel B), and expected heterozygosity computed at the station level (panel C). Color represented the five regions. Expected heterozygosity is presented as box plot and observed values, and differences between regions were tested using one-way ANOVA ($F(4,49) = 9.19$, $p < 0.001$) and Tukey HSD. Stations SFA-0-0, SFA-7-7 and WAZ-3 were excluded from the station level analysis (panel B) given their low sampling size (N < 10). See Fig. 2 for results using the neutral and outlier SNP panel.

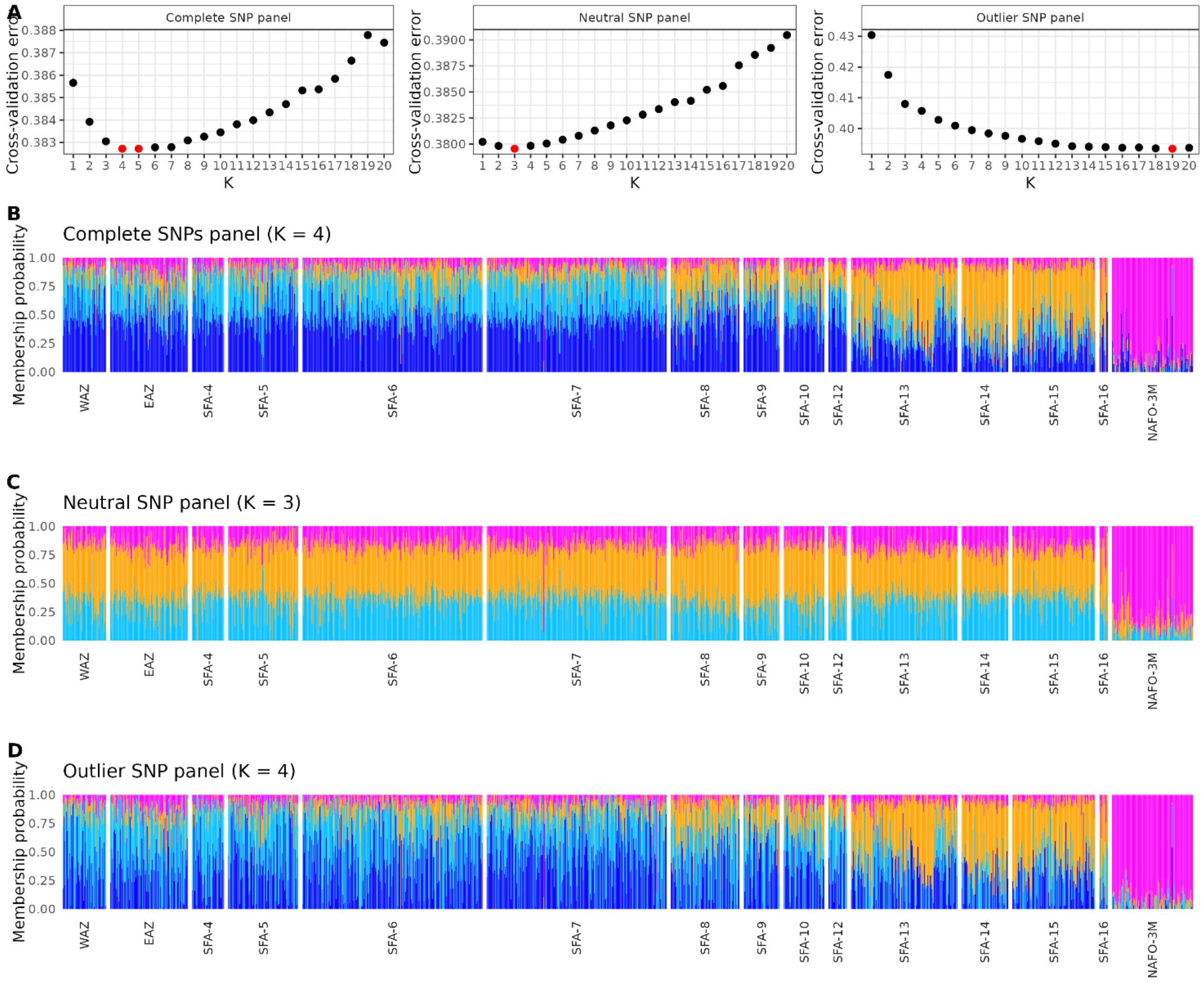


**Figure S7 Admixture results for *P. borealis*, with A) cross-validation tests using the complete, neutral or outlier SNP panel, and membership probability results for B) the complete (K = 4), C) the neutral (K = 3), and D) the outlier SNP panel.** Lowest cross-validation errors observed in panel A are colored in red. In panel B and C, colors represent the inferred clusters for each individual. The membership probability presented for the complete and the neutral SNP panels represented the most probable K, while for the outlier SNP panel, K = 4 was chosen to facilitate the interpretation (no others meaningful group was detected at K > 4).

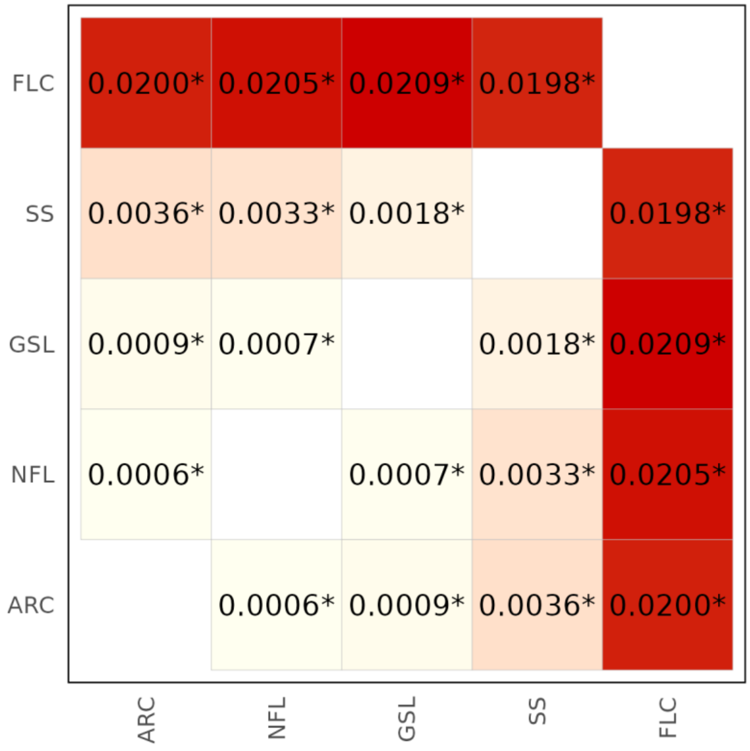


**Figure S8** ***$F_{ST}$* computed among the five regions**. Color gradient reflects $F_{ST}$ values and asterisks (*) represent *P*-value < 0.05.

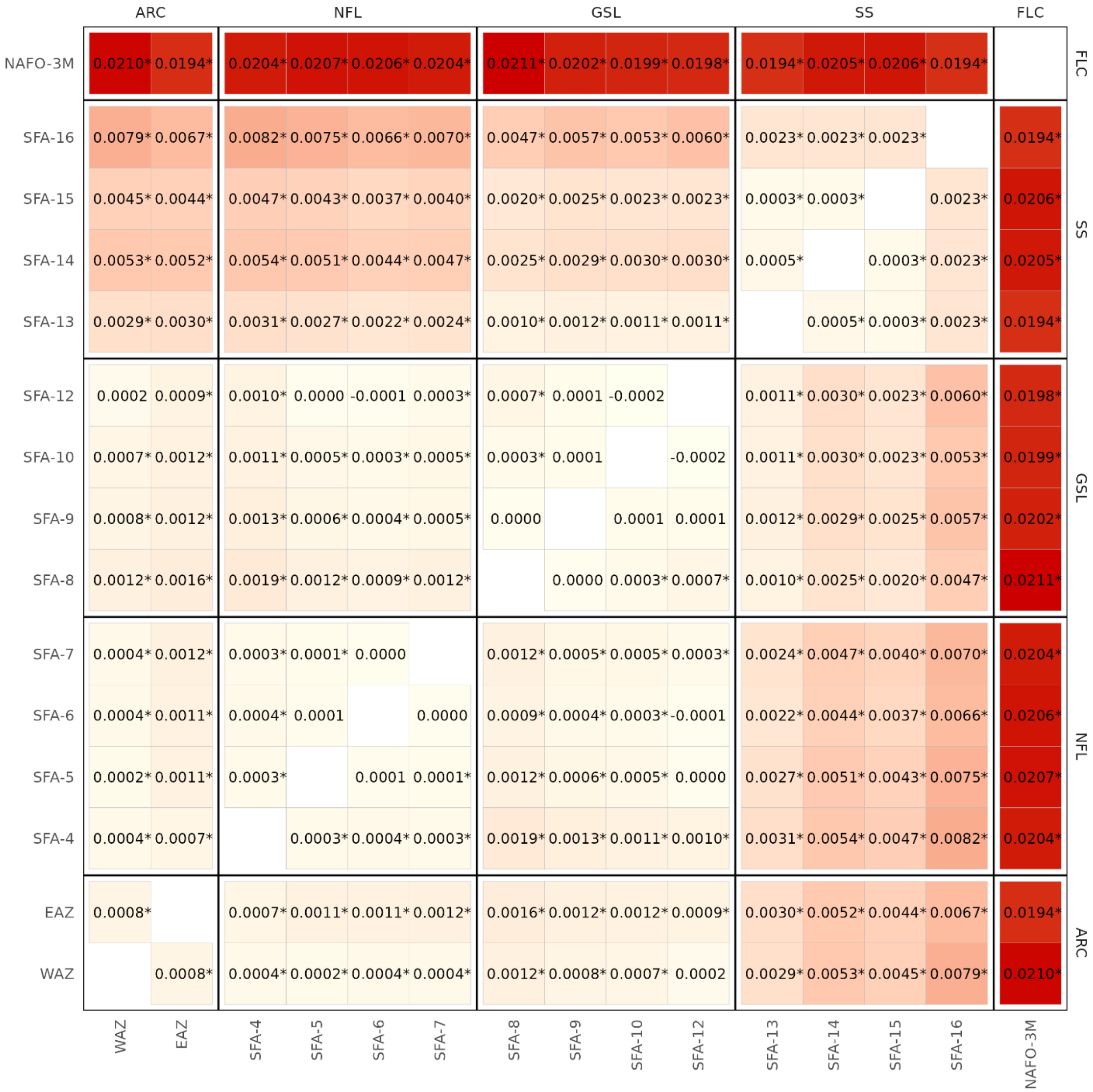


**Figure S9** ***$F_{ST}$* computed among the 15 management areas (WAZ, EAZ, SFA-4 to 16, NAFO-3M).** Color gradient reflects $F_{ST}$ values and asterisks (*) represent *P*-value < 0.05. Management area SFA-0 was excluded given its low sampling size (N = 2).

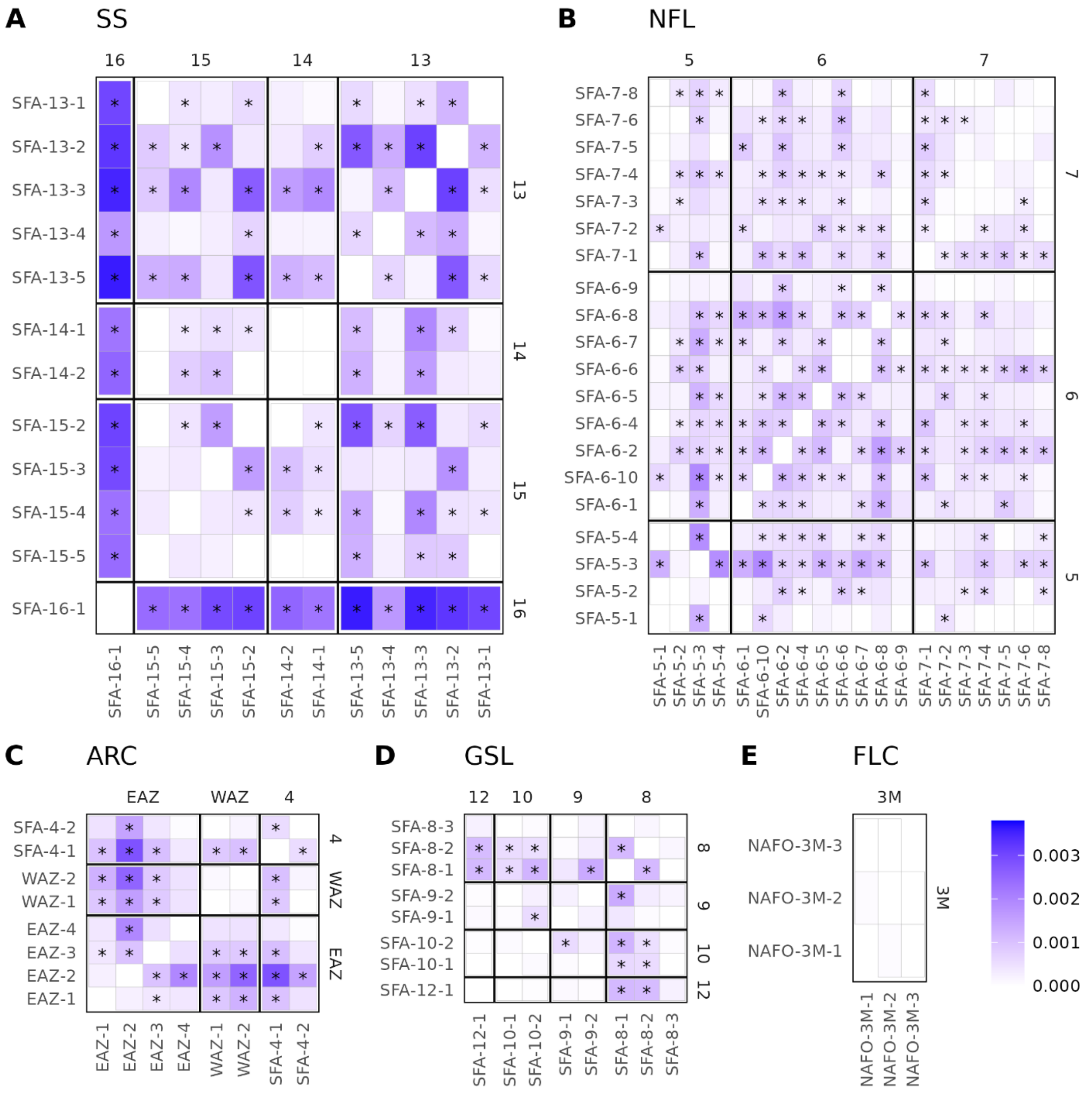


**Figure S10** ***$F_{ST}$* computed among stations within the five regions, A) Scotian Shelf (SS), B) Newfoundland and Labrador (NFL), C) Arctic (AR), D), Gulf of St. Lawrence (GSL) and E) Flemish Cap (FLC).** Color gradient reflects $F_{ST}$ values and asterisks (*) represent *P*-value < 0.05. Stations SFA-0-0, SFA-7-7 and WAZ-3 were excluded given their low sampling size (N < 10).

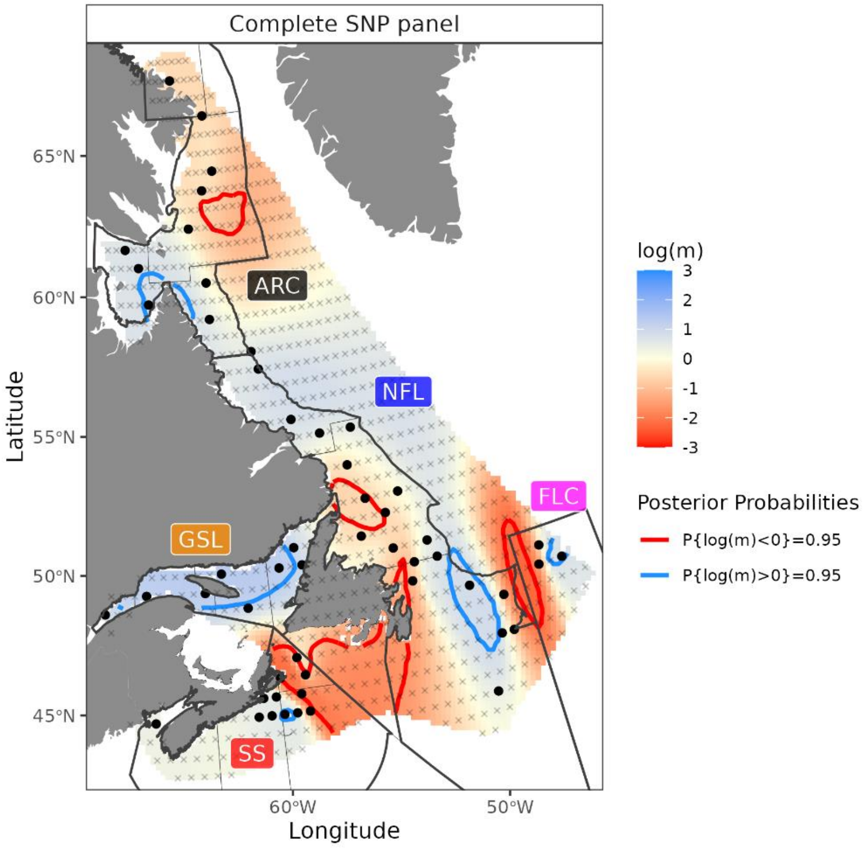


**Figure S11 Estimated effective migration surface, on the log10 scale and following mean centering for all SNPs, using results from 3 chains**. Red areas represented zones of restricted effective migration (i.e., barriers), while blue areas represented zones where exchanges are facilitated (i.e., corridors). Zone of posterior probability P(m ≠ 0 | D) > 95% are delimited by plain color lines. Small gray crosses represented the spatial distribution of demes used (n = 800), while black dots show the location of stations on deme distribution.

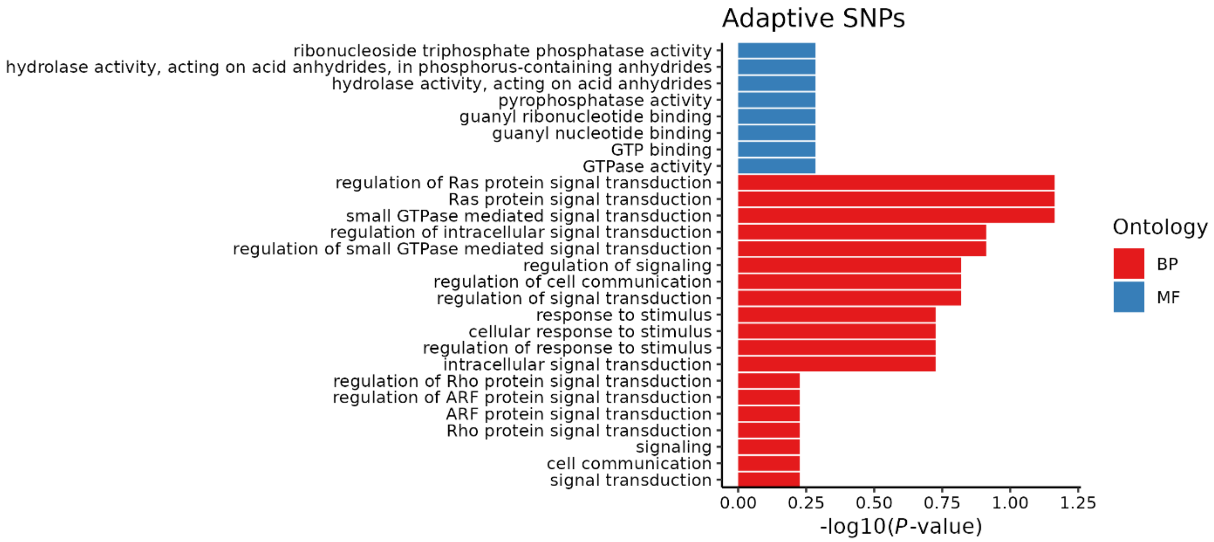


**Figure S12 Functions of transcripts associated to SNPs**. Go enrichment analysis for transcripts associated to adaptive SNPs. BP = Biological Process, MF = Molecular Function.

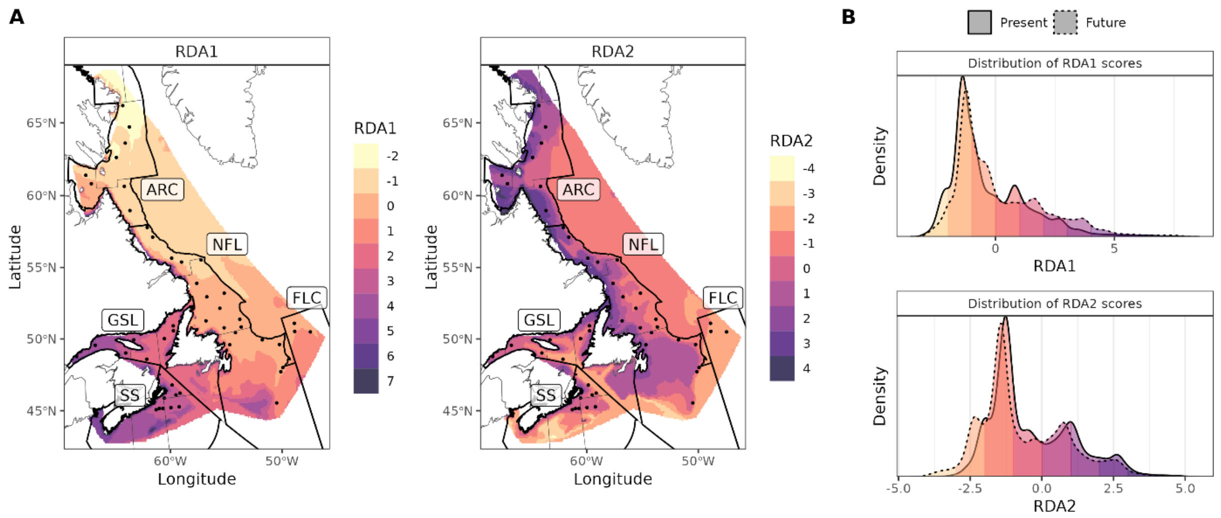


**Figure S13 A) Spatial projection of the adaptive index into the study areas and B) density of RDA scores under present and future environmental condition for RDA1 and RDA2.** Black dots (panel A) represent the stations. See Fig. 4 for projection under current environmental conditions.
